# Molecular analysis of the interactions between phages and the bacterial host *Klebsiella pneumoniae*

**DOI:** 10.1101/2022.09.12.507515

**Authors:** Inés Bleriot, Lucia Blasco, Olga Pacios, Laura Fernández-García, María López, Concha Ortiz-Cartagena, Antonio Barrio-Pujante, Felipe Fernández Cuenca, Álvaro Pascual, Luis Martínez-Martínez, Jesús Oteo-Iglesias, María Tomás

## Abstract

Lytic phages are currently considered among the best options for treating infections caused by multi-drug resistant pathogens. Phages have some advantages over conventional antibiotics. For example, phages acquire modifications in accordance with their environment, and thus with the bacteria present, which has led to the co-evolution of both types of organism. Therefore, both phages and bacteria have acquired resistance mechanisms for protection. In this context, the aims of the present study were to analyze the proteins isolated from twenty-one novel lytic phages of *Klebsiella pneumoniae* in search of defence mechanisms against bacteria and also to determine the infective capacity of the phages. A proteomic study was also conducted to investigate the defence mechanisms of two clinical isolates of *Klebsiella pneumoniae* infected by phages. For this purpose, the twenty-one lytic phages were sequenced and *de novo* assembled using the Illumina-Miseq system and Spades V.3.15.2 respectively. Gene annotation was performed with Patric, Blast, Hhmer and Hhpred tools. The evolutionary relationships between phages were determined by RaxML. The host-range was determined in a collection of forty-seven clinical isolates of *K. pneumoniae*, revealing the variable infectivity capacity of the phages. Genome sequencing showed that all of the phages were lytic phages belonging to the family *Caudovirales*. The size and GC content of the phages ranged from 39,371 to 178,532 bp and from 41.72 % to 53.76 %, respectively. Phage sequence analysis revealed that the proteins were organized in functional modules within the genome. Although most of the proteins have unknown functions, multiple proteins were associated with defence mechanisms against bacteria, including the restriction-modification (RM) system, the toxin-antitoxin (TA) system, evasion of DNA degradation, blocking of host RM, the orphan CRISPR-Cas system and the anti-CRISPR system. Proteomic study of the phage-host interactions (i.e. between isolates K3574 and K3320, which have intact CRISPR-Cas systems, and phages vB_KpnS-VAC35 and vB_KpnM-VAC36, respectively) revealed the presence of several defence mechanisms against phage infection (prophage, plasmid, defence/virulence/resistance and oxidative stress proteins) in the bacteria, and of the Acr candidate (anti-CRISPR protein) in the phages.

**IMPORTANCE:** Phages, viral parasites of bacteria, have long protected the Earth’s biosphere against bacterial overgrowth and could now help in the fight against antimicrobial resistance. However, researchers, including microbiologists and infectious disease specialists, require more knowledge about the interactions between phages and their bacterial hosts and about the defence mechanisms in both viruses and bacteria. In this study, we analyzed the molecular mechanisms of viral and bacterial defence in phages infecting clinical isolates of *Klebsiella pneumoniae*. Viral defence mechanisms included RM system evasion, the Toxin-Antitoxin system, DNA degradation evasion, blocking of host RM and resistance to the abortive infection system (Abi), anti-CRISPR and CRISPR-Cas systems. Regarding bacterial defence mechanisms, proteomic analysis revealed overexpression of proteins involved in the prophage (FtsH protease modulator), plasmid (cupin phosphomannose isomerase protein), defence/virulence/resistance (porins, efflux pumps, LPS, pili elements, quorum network proteins, TA systems and methyltransferases), oxidative stress mechanisms and Acr candidates (anti-CRISPR protein). The findings reveal some important molecular mechanisms involved in the phage-host bacterial interactions; however, further study in this field is required to improve the efficacy of phage therapy.

## INTRODUCTION

Bacteriophages, or phages, are natural predators of bacteria. Phages are the most abundant and ubiquitous biological entities on Earth, accounting for an estimated total of 10^31^ viral particles (1, 2). In the current context of increasing antibiotic resistance, the emergence of alternative therapies is welcome. Thus, lytic phages are currently considered one of the best options for treating infections caused by multidrug-resistant (MDR) bacteria (3, 4), as demonstrated in clinical trials conducted to date (5–10). In general, phage therapy has some advantages over the use of conventional antibiotics, such as low toxicity and high host specificity (3, 11). These characteristics enable phages to target the pathogen responsible for the infection to be treated, thus preserving the commensal or mutualistic bacteria that make up the microbiota, whose role in human health we are only beginning to understand. In addition, phages are usually easier to administer, as they do not require repeated administration, as is generally required with antibiotics. This is because phages divide at the site of infection and can therefore remain in the human body for relatively long periods of time. Another characteristic of phages that makes them good candidates for therapy is their ability to adapt to changes in the bacterial host, which has resulted from the coevolution of both types of organisms (12).

Bacteria have developed mechanisms to prevent phage infection at almost all stages of the viral replication cycle (13, 14). Firstly, bacteria can prevent phage attachment by mutating or altering their surface receptors, by producing inhibitors that outcompete the phage for receptors, or by producing polysaccharides that physically mask phage receptors (13). In order to prevent injection of phage DNA into the cytoplasm, the bacteria then use the superinfection exclusion (Sie) system, which is characterized by proteins that block the entry of phage DNA into host cells (15). If the phage nevertheless manages to enter the cell, the bacteria employ other defence systems such as the abortive infection system (Abi) to interrupt phage development at any stage (replication, transcription or translation) (16), the bacteriophage exclusion system (BREX) (17) or the defence island system associated with restriction-modification (DISARM) (18) to interrupt replication. Phages can also use the Toxin-Antitoxin (TA) systems, characterized by adjacent genes, generally consisting of two components: a stable toxin and an unstable antitoxin. The unstable component is degraded under stress conditions by the protease system, leading to toxin activation and often resulting in reduced bacterial metabolism and phage inhibition (19, 20). Bacteria can also employ other types of systems to cleave phage DNA, such as the restriction-modification (RM) system, which is characterized by an endonuclease protein that protects the bacterial cell by cutting foreign DNA at specific points, and a cognate DNA methylase, which modifies and protects the host DNA (21). They can also employ the clustered regularly interspaced short palindromic repeats–CRISPR-associated proteins (CRISPR-Cas) system, an adaptive immune system (22) characterized by the acquisition of spacer sequences, which are small fragments of foreign nucleic acids of phage or foreign DNA, between the repeats of the CRISPR locus (23).

Phages have, in turn, developed counterstrategies to evade bacterial defence mechanisms (24). For example, for successful adsorption, phages can modify their receptor binding proteins (RBPs) by acquiring mutations to obtain new receptors (25). In turn, they are also able to acquire enzymes such as depolymerase to access masked receptors (26, 27) in a way that allows them to interact with a surface component expressed by the host at that time (28). However, when a phage genome manages to enter the cell, it can still face the myriad intracellular antiviral barriers described above. Phages can respond to these by promoting the mutation of specific genes to prevent activation of the bacterial Abi system (29). They can also evade bacterial RM systems by reducing the number of restriction sites in their genome (30), modifying bases in their genome (31), co-injecting protein (for instance, DarA and DarB in the phage P1) with the genome to bind directly to the phage DNA and mask restriction sites (32), stimulating the action of modification enzymes and degrading an RM cofactor. Phages can also sequester host antitoxins via a protein that probably inhibits Lon protease activity, to avoid the deleterious action of the toxins of the TA systems (33). To circumvent the effect of bacterial TA systems, one phage, T4, encodes its own antitoxin protein (Dmd) that functionally replaces the unstable antitoxin of the host, thereby promoting phage propagation (34). Finally, phages have developed mechanisms to evade the bacterial CRISPR-Cas system. For instance, they can evade CRISPR through a single-nucleotide substitution or a complete deletion in the protospacer region or in the conserved protospacer-adjacent motif (PAM) (35). Phages have also developed anti-CRISPR systems, which basically consist of Acr proteins (typically small proteins of 80-150 aa) that inhibit bacterial CRISPR-Cas activity by binding directly to, and thus inactivating, the Cas protein, so that phages can successfully replicate in the bacterial host (36).

A better understanding of phage-host interaction could lead to the development of more successful therapeutic applications for phages. In this context, the aims of the present study were to analyze the proteins isolated from twenty-one novel lytic phages of *K. pneumoniae* in search of defence mechanisms against bacteria and also to determine the infectivity capacity of the phages. A further aim was to investigate the defence mechanisms of bacteria in response to phage infection.

## MATERIAL AND METHODS

### Bacterial strains

A collection of forty-seven clinical isolates of carbapenemase-producing *K. pneumoniae* obtained from the Virgen Macarena University Hospital (Seville, Spain) and the National Centers for Microbiology (CNM; Carlos III Health Institute, Spain) was used in this study (Table 1). The sequence type (ST) and the capsular type (K) of each strain were determined using the methods available on the Pasteur Institute website (http://bigsdn.web.pasteur.fr/Klebsiella, accessed between 2018 and present) and the Kaptive website (https://Kaptive-web-erc.monash.edu, accessed in April 2020), respectively. All strains were grown in Luria-Bertani (LB) medium (0.5 % NaCl, 0.5 % yeast extract, 1 % tryptone).

**Table 1.**
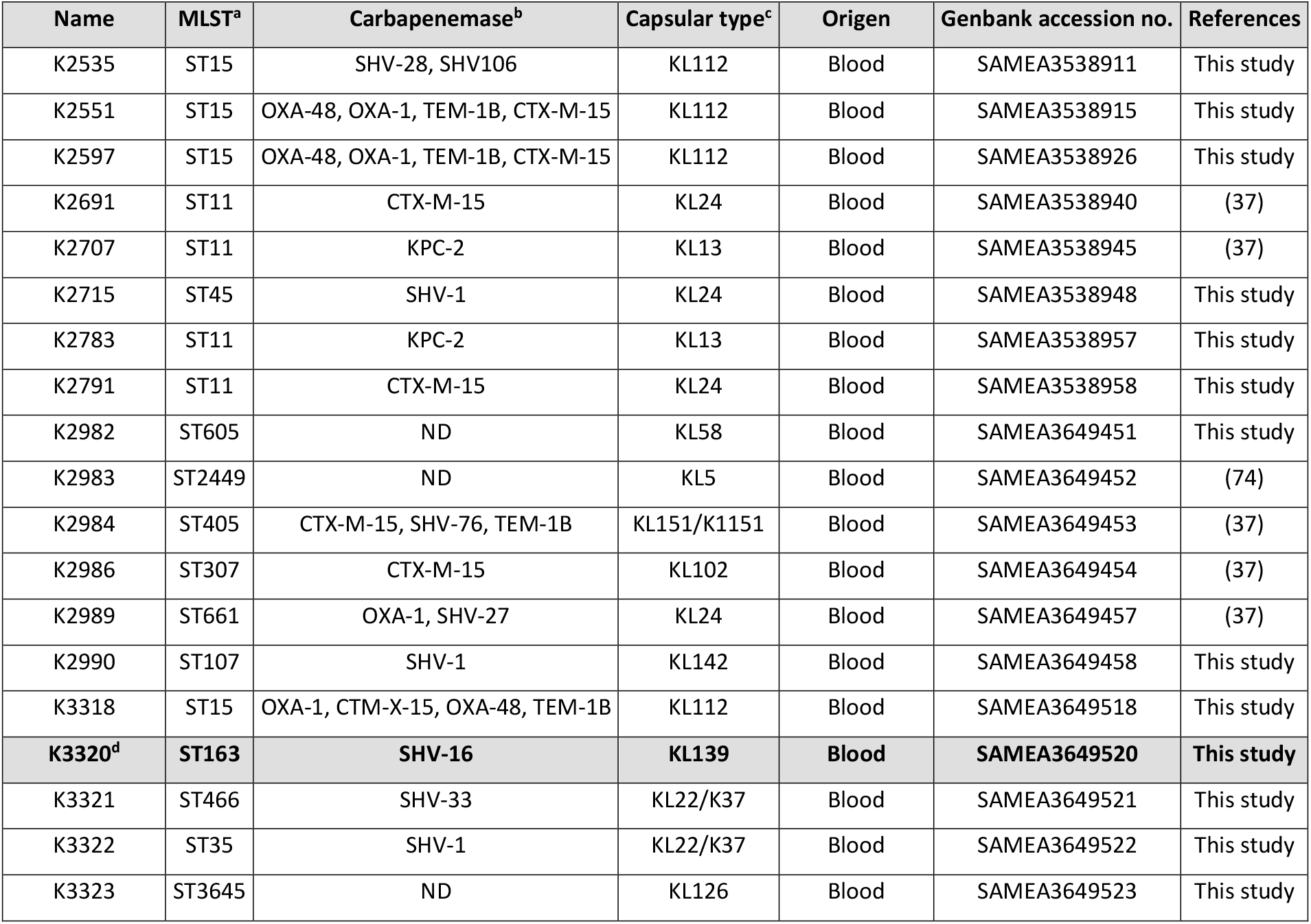
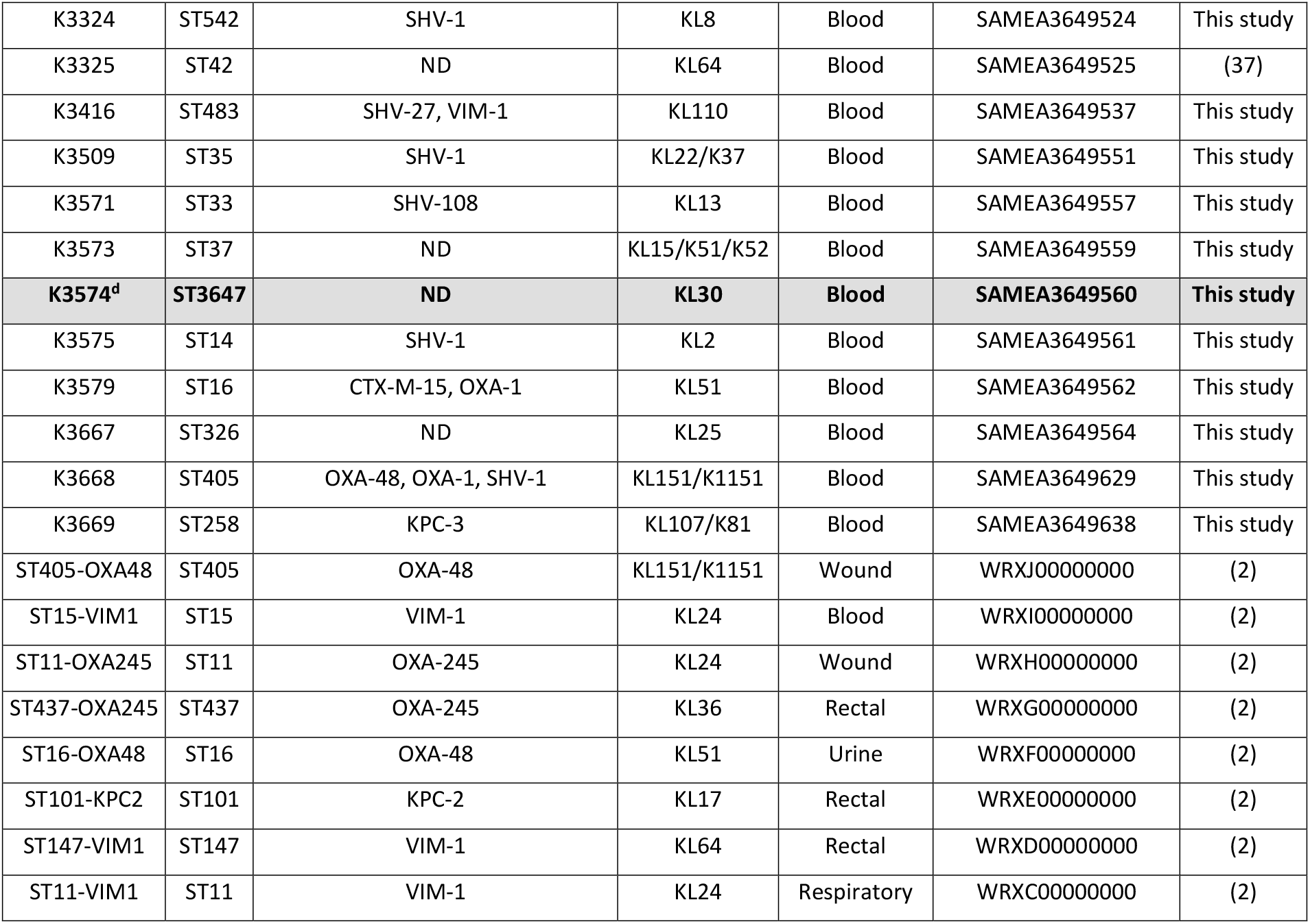
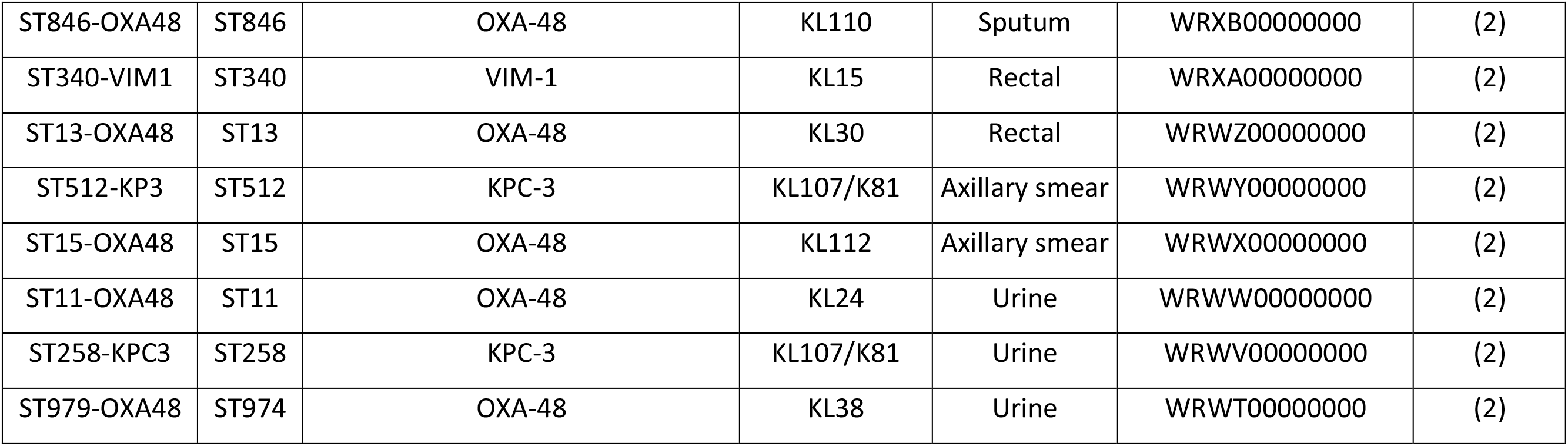
Characteristic of the collection of the forty-seven clinical strains of *K. pneumoniae*.^a^ MSLT, determined by http://bigsdb.pasteur.fr/Klebsiella.html, ^b^ Carbapenemase, determined by https://cge.cbs.dtu.dk/services/ResFinder/, ^c^ Capsular type, determined by https://kaptive-web.erc.monash.edu. ^d^ In bold, the clinical strains selected for phage interaction studies.

### Isolation and purification of lytic phages

Ten new lytic phages isolated from sewage water samples and twelve lytic phages previously isolated by our research group (20, 37, 38) were used in this study. Briefly, 50 mL samples of water were collected near sewage plants and held at room temperature until processing in the laboratory. Once in the laboratory, the samples were vortexed and centrifuged at 4000 × g for 10 min. The supernatant was recovered and filtered through membranes of pore sizes 0.45 μm and 0.22 μm, to remove debris. One-mL aliquots of the filtered samples were then added to 500 μL of the natural host *K. pneumoniae* (Table 2) in 4 mL of soft agar (0.5 % NaCl, 1 % tryptone and 04 % agar; supplemented or not with 1 mM CaCl_2_) and poured onto TA agar plate (0.5 % NaCl, 1 % tryptone and 1.5 % agar; supplemented or not with 1 mM CaCl_2_) (i.e. the double-layer agar technique). The plates were incubated at 37 ºC. Isolated plaques of different morphology were then recovered by picking with a micropipette and were stored at – 80ºC. Two additional plaque assays and plaque picking steps were performed to check and purify the isolated plaques.

**Table 2.** Characteristics of the twenty-one lytic phages of *K. pneumoniae*. In bold, the clinical strains selected for phage interaction studies.

| Phage | Natural host | Acc. Number | Family | Genus | Genome size (bp) | G+C % | References |
| --- | --- | --- | --- | --- | --- | --- | --- |
| vB_KpnP-VAC1 | ATCC10031 | SAMN19773206 | *Autographiviridae* | *Teetrevirus* | 39,371 | 50.75 % | (20) |
| vB_KpnS-VAC2 | ATCC10031 | SAMN19773207 | *Drexelviriidae* | *Webervirus* | 51,784 | 47.86 % | (20) |
| vB_KpnS-VAC4 | ATCC10031 | SAMN19773215 | *Drexelviriidae* | *Webervirus* | 45,558 | 51.11 % | (20) |
| vB_KpnS-VAC5 | ATCC10031 | SAMN19773216 | *Drexelviriidae* | *Webervirus* | 49,636 | 50.46 % | (20) |
| vB_KpnS-VAC6 | ATCC10031 | SAMN19773219 | *Drexelviriidae* | *Webervirus* | 51,554 | 51.63 % | (20) |
| vB_KpnS-VAC7 | ATCC10031 | SAMN19773224 | *Drexelviriidae* | *Webervirus* | 49,684 | 51.25 % | (20) |
| vB_KpnS-VAC8 | ATCC10031 | SAMN19773221 | *Drexelviriidae* | *Webervirus* | 48,933 | 50.52 % | (20) |
| vB_KpnS-VAC10 | ATCC10031 | SAMN19773232 | *Drexelviriidae* | *Webervirus* | 48,935 | 50.65 % | (20) |
| vB_KpnS-VAC11 | ATCC10031 | SAMN19773540 | *Drexelviriidae* | *Webervirus* | 48,826 | 50.83 % | (20) |
| vB_KpnM-VAC13 | ATCC10031 | SAMN22059222 | *Myoviridae* | *Slopekvirus* | 174,826 | 41.93 % | (37) |
| vB_KpnM-VAC25 | K3579 | SAMN20298869 | *Autographiviridae* | *Drulisivirus* | 43,777 | 53.76 % | This study |
| **vB_KpnS-VAC35** | **K3574** | **SAMN20298871** | ***Ackermannviridae*** | ***Taipeivirus*** | **112,862** | **45.44 %** | **This study** |
| **vB_KpnM-VAC36** | **K3573** | **SAMN20298872** | ***Myoviridae*** | ***Slopekvirus*** | **169,970** | **40.90 %** | **This study** |
| vB_KpnS-VAC51 | K3325 | SAMN23489160 | *Ackermannviridae* | *Taipeivirus* | 113,149 | 45.40 % | This study |
| vB_KpnM-VAC66 | K3320 | SAMN22059211 | *Myoviridae* | *Slopekvirus* | 178,532 | 41.72 % | (38) |
| vB_KpnS-VAC70 | K3318 | SAMN20298916 | *Drexelviriidae* | *Webervirus* | 49,631 | 50.70 % | This study |
| vB_KpnP-VAC71 | K3318 | SAMN20298917 | *Autographiviridae* | *Przondovirus* | 40,388 | 52.98 % | This study |
| vB_KpnS-VAC110 | K2691 | SAMN24377650 | *Drexelviriidae* | *Webervirus* | 45,195 | 50.44% | This study |
| vB_KpnS-VAC111 | K2691 | SAMN20298918 | *Drexelviriidae* | *Webervirus* | 50,559 | 50.5 % | This study |
| vB_KpnS-VAC112 | K2691 | SAMN20298919 | *Drexelviriidae* | *Webervirus* | 49,068 | 50.20 % | This study |
| vB_KpnS-VAC113 | K2691 | SAMN20298921 | *Drexelviriidae* | *Webervirus* | 49,568 | 50.46 % | This study |

### Propagation of phage and transmission electron microscopy

Plaque-purified phages were amplified in LB liquid media (supplemented or not with 1 mM CaCl_2_, depending on phage), with shaking (180 rpm) at 37 ºC, by infecting an early logarithmic growth phase (OD_600nm_ = 0.3 – 0.4) of the natural host of each phage (Table 2). After lysis, i.e. when the culture appeared clear, bacterial debris was removed by centrifugation (4302 × g 10 min) and the remaining suspension was filtered through membranes of pore sizes 0.45 μm and 0.22 μm. Finally, the supernatants were serially diluted in SM buffer (0.1 M NaCl, 10 mM MgSO_4_, 20 mM Tris-HCL pH 7.5) and seeded by the double-layer agar method. The ten new lytic phage solutions were negatively stained with 1 % aqueous uranyl acetate before being analyzed by transmission electron microscopy (TEM) in a JEOL JEM-1011 electron microscope.

### Phage DNA extraction and whole genome sequencing (WSG)

The phage DNA of the ten new lytic phages was isolated from the strains with the phenol:chloroform method following the phagehunting protocol (http://phagesdb.org/media/workflow/protocols/pdfs/PCI_SDS_DNA_extraction_2.2013.pdf, accessed on 1 February 2021). DNA concentrations and quality were measured in a Nanodrop ND-10000 spectrophotometer (NanoDrop Technologies, Waltham, MA, USA) and Qubit fluorometer (Thermo Fisher Scientific, USA). Genomic libraries were then prepared using the Nextera XT Library prep kit (Illumina), following the manufacturer’s instructions. The distribution of fragment lengths was checked in an Agilent 2100 Bioanalyser, with the Agilent Hight sensitivity DNA kit. Libraries were purified using the Mag-Bind RXNPure plus magnetic beads (Omega Biotek) and finally, the pool was sequenced in Miseq platform (Illumina Inc, USA). The quality of the FASTQ file was checked using the software FastQC (39) and summarized using MultiQC (40). Sequences of 300 bp paired-end reads of each isolate were *“de novo”* assembled using Spades V.3.15.2 (41).

### Phage genome annotation

#### Defence mechanisms

All assemblies were initially annotated by sequence homology using Patric 3.6.9 (http://patricbrc.org, accessed on 22 February 2021) and were then manually refined using Blastx (http://blast.ncbi.nlm.nih.gov, accessed between August and October 2021) and Hhmer (http://hmmer.org, accessed between August and October 2021), as well as the Hhpred tool (https://toolkit.tuebingen.mpg.de/tools/hhpred, accessed between August and October 2021), which predict functions through protein structure. In addition, to search for phage defence mechanisms against bacteria, the CRISPR Miner 2 (http://www.microbiome-bigdata.com/CRISPRminer2/index/, accessed in March 2022) and PADLOC (https://padloc.otago.ac.nz/padloc/, accessed in March 2022) tools were used to search for possible CRISPR-cas systems, as well as the AcrDB tool (https://bcb.unl.edu/AcrFinder/, accessed on October 2021) to search for possible anti-CRIPSR-cas systems with the defect parameter of the website (Aca e-value: 0.01, Aca identity %: 30. Aca coverage: 0.8, Maximum intergenic distance between genes [bp]: 150; Operon up/down-stream range for MGE-Prophage search [no. of genes]: 10). Finally, the family and genus of the different phages were determined by sequence homology with the phage sequences available in the NCBI database. Complete genome sequences were included in the GenBank Bioproject PRJNA739095 (http://www.ncbi.nlm.giv/bioproject/739095).

#### Phage phylogenetic analysis and genome comparison

Phylogenetic analysis of the twenty-one phages was performed using the nucleotide sequence of the large terminase subunit of each phage. Alignment was first performed with MAFF server (https://mafft.cbrc.jp/alignment/server/index.html, accessed on 3 January 2022) and a phylogenetic tree was then constructed using RAxMLHPC-PTHREADS-AVX2 version 8.2.12 (42) under the GTRGAMMA model and 100 bootstrap replicates. A graphical representation of the comparison of all phage genomes was then constructed with the VipTree website (https://www.genome.jp/viptree/, accessed in June 2022) according to the previously established phylogenetic relationship.

### Host-range assay

The phage host spectrum was tested by the spot test technique (43), in a collection of forty-seven clinical strains of *K. pneumoniae*. Briefly, 200 μL of an overnight culture was mixed with 4 mL soft agar and poured on TA agar plates. Once the soft medium solidified, 15 μL drops of high titre phages were added to the plates. For each isolate, a negative control consisting of SM buffer was included in each plate. All determinations were made in triplicate. The criteria used to determine the phage infectivity were lack of spots (no infection), presence of clear spots (infection) and presence of turbid spots (low infection or resistance).

### Study of bacteria genome of the K3574 and K3320 clinical isolates

The genome of the clinical isolates of *K. pneumoniae* K3574 (SAMEA3649560) and K3320 (SAMEA3649520) was examined to search CRISPR-Cas systems, by using the CRISPR Miner 2 (http://www.microbiome-bigdata.com/CRISPRminer2/index/, accessed in March 2022) and the PADLOC tool (https://padloc.otago.ac.nz/padloc/, accessed in March 2022). In addition, the genomic annotation of the Rastserver was also studied in order to check and validate the results obtained with other tools (https://rast.nmpdr.org, accessed in August 2022). Plasmids were then searched for using the plasmidfinder v2.0.1 (20-07-01) (https://cge.food.dtu.dk/services/PlasmidFinder/, accessed in July 2022), RM system using the Restriction-ModificationFinder v.1.1 (accessed in June 2015) (https://cge.food.dtu.dk/services/Restriction-ModificationFinder/history.php, accessed in July 2022) and prophages using the Phaster tools (http://phaster.ca, accessed in July 2022).

### Characterization of phages vB_KpnS-VAC35 and vB_KpnM-VAC36

#### Phage adsorption

Adsorption of phages vB_KpnS-VAC35 and vB_KpnM-VAC36 to the bacterial surface receptors of clinical strains K3574 and K3320, respectively, was determined from the adsorption curve (44). Briefly, overnight cultures of *K. pneumoniae* K3574 and K3320 were diluted 1:100 in LB and incubated at 37 ºC at 180 rpm, until a cell count of 10^8^ CFU/mL was reached. At this point, the cultures were held at room temperature without shaking and were infected with a phage suspension at multiplicity of infection (MOI) of 0.01. Every 2 minutes, 1 mL of culture was removed and placed in contact with 1 % of chloroform. The samples were then centrifuged for 2 min at 12000 × g to sediment the cell debris and adsorbed phage. The supernatants were serially diluted in SM buffer for subsequent plating on a double agar plate with the corresponding host plating (K3574 and K3320, respectively). The number of phages mixed with bacterial host cells at time 0 was considered 100 % free of phages. The adsorption curve analysis was performed in triplicate.

#### One-step growth curve assay

A one-step growth curve of phages vB_KpnS-VAC35 and vB_KpnM-VAC36 was constructed in clinical strains K3574 and K3320, respectively, to determine the latent period (L) and the burst size (B). The latent period was defined as the interval between adsorption of the phages to the bacterial cells and the release of phage progeny. The phage burst size was defined as the number of viral particles released in each cycle of infection per bacteria cells. For this purpose, an overnight culture of the strains was diluted 1:100 in LB and incubated at 37 ºC at 180 rpm, until a cell count of 10^8^ CFU/mL was reached. At this point, cultures were infected with a phage suspension at an MOI of 0.01 and held at room temperature for the respective adsorption times (5 and 2 min). The cultures were then washed twice by centrifugation at 6000 × g for 10 min in order to remove the free phages. The pellet was then resuspended in 1 mL of LB, and 25 μL of bacterial mixture was added to 25 mL of LB (time 0). Dilutions were made in SM buffer and subsequently seeded on double agar plates for subsequent quantification. The one-step growth curve analysis was performed in triplicate.

#### Phage kill curve assay in liquid medium

Killing curves were constructed from the selected isolates K3574 and K3320, in accordance with the presence of intact CRISPR-cas system in their genome. Phages vB_KpnS-VAC35 and vB_KpnM-VAC36 were used to monitor the infection of the strain by optical density measured at a wavelength of 600 nm (OD_600nm_) and counts of CFU/mL and PFU/mL. For this purpose, an overnight culture of the selected strains was diluted 1:100 in LB and incubated at 37 ºC at 180 rpm, until the early logarithmic phase was reached (OD_600nm_ = 0.3 – 0.4). At this point, cultures were infected with phages at an MOI of 1. The OD_600nm_, the number of CFU/mL and PFU/mL were determined every 30 min for 3 h. In all cases, the control was the strain without phage infection. All analyses were performed in triplicate.

### NanoUHPLC-Tims-QTOF proteomic analysis: interaction between phages (vB_KpnS-VAC35 and vB_KpnM-VAC36) and clinical strains (K3574 and K3320)

NanoUHPLC-Tims-QTOF analysis was performed for quantitative study of the protein profile of strain K3574 and K3320 with and without infection with phages vB_KpnS-VAC35 and vB_KpnM-VAC36. The samples were first prepared and overnight culture of strains was diluted 1:100 in 25 mL LB, and incubated at 37 ºC at 180 rpm, until the cultures reached an early logarithmic phase of growth (OD_600nm_ = 0.3 - 0.4). The cultures were then infected with phages at an MOI of 1. After 1 h, the cultures of the strains were harvested by centrifugation at 4302 × g for 20 min at 4 ºC. The pellets were then stored at – 80 ºC to facilitate cell disruption. The next day, the pellet was resuspended in PBS and sonicated. Finally, the sonicated pellets were centrifuged at 4302 × g for 20 min at 4 ºC, and the flow-through, i.e. crude extract, was analyzed by LC-MS and NanoUHPLC-Tims-QTOF. The equipment used for this purpose was a TimsTof pro mass spectrophotometer (Bruker), a nanoESI source (CaptiveSpray), a time-QTOF analyser and a nanoELUTE chromatograph (Bruker). Sample preparation was carried out by tryptic digestion in solution with reduction-alkylation followed by Ziptip desalting. Data were acquired in nanoESI positive ionization mode, Scan PASEF-MSMS mode and CID fragmentation mode, with an acquisition range of 100-1700 m/z. The products were separated on a Reprosil C18 column (150 × 0.075 mm, 1.9 μm and 120 Å) (Bruker) at 50 ºC, with an injection volume of 2 μL. The mobile phases consisted of (A) 0.1 % H_2_O/formic acid and (B) 0.1 % acetonitrile/formic acid. The flow rate was 0.4 μL/min, and the gradient programme was as follows: 11 % B (0-5 min), 16 % B (5-10 min), 35 % B (10-16 min), 95 % B (16-18 min) and 95 % B (18-20 min). Finally, different software was used for data acquisition: Compass Hystar 5.1 (Bruker) and TimsControl (Brunker) and TimsControl (Brunker), Data analysis (Bruker) and PEAKS studio (Bioinformatics solutions).

## RESULTS

### Isolation, propagation and electron microscopic analysis of phages

The twenty-one phages used in this study, named according to accepted practices (45) (Table 2), were obtained from wastewater samples. However, to facilitate reading this paper, we will incorporate the data previously found in these studies in order to enable comparison between the phages. Thus, TEM studies revealed that fifteen phages belonged to the *Siphoviridae* family, characterized by a long, flexible tail (vB_KpnS-VAC2, vB_KpnS-VAC4, vB_KpnS-VAC5, vB_KpnS-VAC6, vB_KpnS-VAC7, vB_KpnS-VAC8, vB_KpnS-VAC10 and vB_KpnS-VAC11 (20), vB_KpnS-VAC35, vB_KpnS-VAC51, vB_KpnS-VAC70, vB_KpnS-VAC110, vB_KpnS-VAC111, vB_KpnS-VAC112 and vB_KpnS-VAC113). Moreover, four (vB_KpnM-VAC13 [37], vB_KpnM-VAC25, vB_KpnM-VAC36 and vB_KpnM-VAC66 [38]) were found to be members of the family *Myoviridae*, which is characterized by an icosahedral capsid and a rigid, contractile tail. Finally, only two phages were identified as members of the family *Podoviridae*, characterized by a small, non-contractile tail (vB_KpnP-VAC1 [20] and vB_KpnP-VAC71) (Figure 1B).

**Figure 1.**
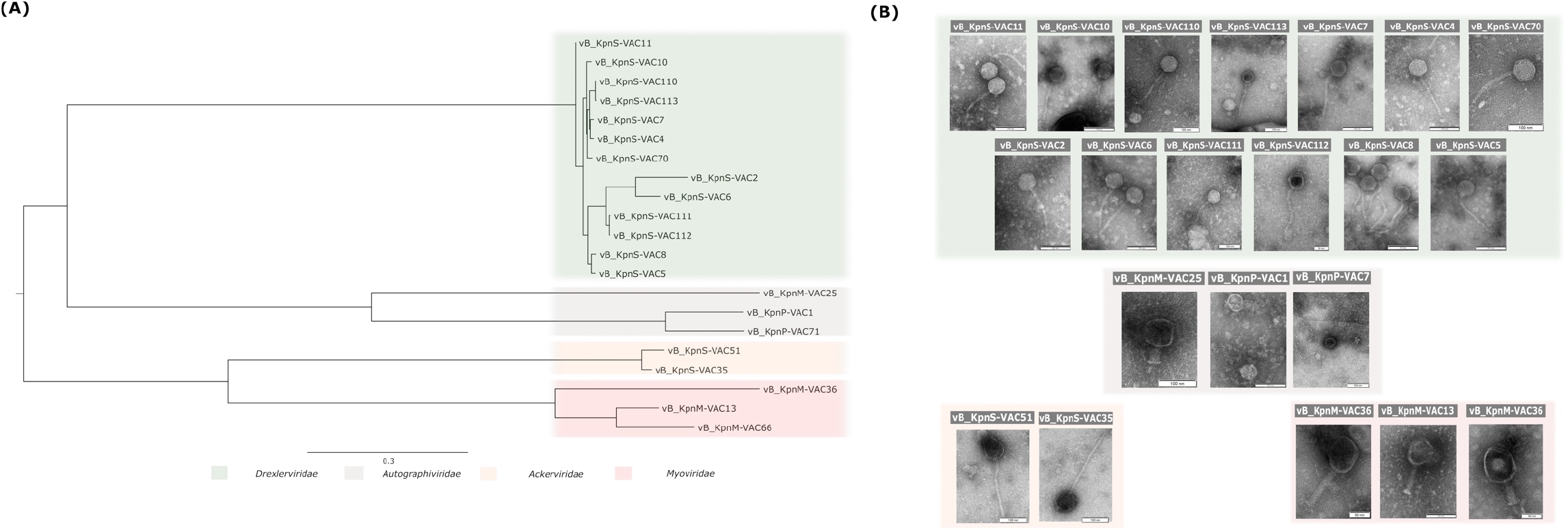
**(A)** Phylogenetic analysis of the twenty-one phages performed with the nucleotide sequence of the large terminase subunit of each phage. **(B)** Transmission electron microscopy images showing the structure of the twenty-one phages under study. All belong to the order *Caudovirales*. vB_KpnP-VAC1 and vB_KpnP-VAC71 are members of the *Podoviridae* family, characterized by a short tail. On the other hand, phages vB_KpnS-VAC2, vB_KpnS-VAC4, vB_KpnS-VAC5, vB_KpnS-VAC6, vB_KpnS-VAC7, vB_KpnS-VAC8, vB_KpnS-VAC10, vB_KpnS-VAC11, vB_KpnS-VAC35, vB_KpnS-VAC70, vB_KpnS-VAC110, vB_KpnS-VAC111, vB_KpnS-VAC112 and vB_KpnS-VAC113 are members of the *Siphoviridae* family, characterized by large, flexible tails. Finally, phages vB_KpnM-VAC13, vB_KpnM-VAC35 and vB_KpnM-VAC66 are members of the *Myoviridae* family, characterized by an icosahedral capsid and a rigid, contractile tail. The scale bar of the TEM represents 50 or 100 nm, depending on the phage.

### Phage genome annotation

#### Phage genome analysis

The phage genome sequencing revealed that all phages under study, available from the Genbank Bioproject PRJNA739095 (http://www.ncbi.nlm.nih.gov/bioproject/739095) (Table 2), were lytic *Caudovirales* phages, i.e. dsDNA tailed phages, lacking lysogenic genes such as integrase, recombinase and excesionase. More specifically, 63.63 % of the phages belonged to the *Drexerviridae* family (14 of 21 phages), 13.64 % belonged to the *Autographiviridae* family (3 of 21 phages), 13.64 % belonged to the *Myoviridae* family (2 of 21 phages) and 9.09 % belonged to the *Ackermannviridae* family (2 of 21 phages). The genomes ranged from 39,371 bp in the case of phage vB_KpnP-VAC1, a member of the family *Autographiviridae* and the genus *Teetrevirus*, to 178,532 bp in the case of phage vB_KpnM-VAC66, a member of the family *Myoviridae*, subfamily *Tevenvirinae* and the genus *Slopekvirus*. The guanine-cytosine content ranged from 40.90 % in the case of vB_KpnM-VAC36 to 53.76 % in the case of vB_KpnM-VAC25. The genomic study revealed that the structure of the phage genomes varied depending on the type of phage. In the case of the members of the *Drexerviridae* (vB_KpnS-VAC2-11, vB_KpnS-VAC70 and vB_KpnS-VAC110-113), Autographiviridae (vB_KpnP-VAC1, vB_KpnP-VAC25 and vB_KpnP-VAC71) and *Ackermannviridae* (vB_KpnS-VAC35 and vB_KpnS-VAC51) families, the genome was organized in functional modules of genes related to structure, packaging, lysis, transcription and regulation. By contrast, for members of the *Myoviridae* (vB_KpnM-VAC13, vB_KpnM-VAC36 and vB_KpnM-VAC66) family, which are “larger phages” (> 100 bp), no lysis-specific blocks were distinguished, and structural and morphogenesis-related proteins were repeated in several blocks throughout the genome. Considering the lysis genes, all phages had endolysins and holins, proteins which are responsible for the degradation of the bacterial cell wall during infection of the host. However, a difference was observed in terms of spanin, a protein involved in the lysis process in Gram-negative hosts, depending on the family to which the phage belongs: the *Drexerviridae* family had a unimolecular spanin (U-spanin), while the *Autographiviridae*, *Ackermannviridae* and *Myoviridae* families had a heterodimer molecule of spanin (I-spanin and O-spanin). Regarding the depolymerase genes, generally related to the tail receptor, five of the phages (23.81 %) had one depolymerase (vB_KpnS-VAC4, vB_KpnS-VAC7, vB_KpnS-VAC10, vB_KpnS-VAC11 and vB_KpnS-VAC70), while two of the phages (9.52 %) (vB_KpnS-VAC2 and vB_KpnS-VAC6) had two different depolymerase genes. It has been observed that the “larger phages” (vB_KpnM-VAC13, vB_KpnS-VAC35, vB_KpnM-VAC36, vB_KpnS-VAC51 and vB_KpnM-VAC66) had numerous tRNA genes (1, 21, 7, 22 and 1 respectively). The presence of HNH homing endonucleases has been observed in the genomes of three phages (vB_KpnP-VAC1, vB_KpnM-VAC13 and vB_KpnM-VAC66). The phage vB_KpnM-VAC66 contains the highest number of HNH homing endonucleases.

#### Genetic defence mechanisms of phages

An in-depth study of the phage genomes also revealed the presence of defence mechanisms (Table 3): thirty-five restriction-modification (RM) system evasion systems located in sixteen phages, six TA systems located in two phages (vB_KpnM-VAC13 and vB_KpnM-VAC66), one DNA degradation evasion located in phage vB_KpnP-VAC1, four blocking RM of host bacteria located in phage vB_KpnM-VAC36, seven genes that confer resistance to the abortive infection (Abi) system of host bacteria located in three phages (vB_KpnM-VAC13, vB_KpnM-VAC36 and vB_KpnM-VAC66) and finally, two possible orphan CRISPR-Cas system was located in the phages vB_KpnS-VAC35 and vB_KpnS-VAC51. In addition, almost all phages possessed a possible anti-CRISPR system, composed by Acr and Aca protein, except for phages vB_KpnS-VAC112 and vB_KpnS-VAC113 (supplementary Table 1). An inhibitor of the TA system (protein ID: QZE51102.1) was also found in the genome of the phage vB_KpnP-VAC1.

**Table 3.**
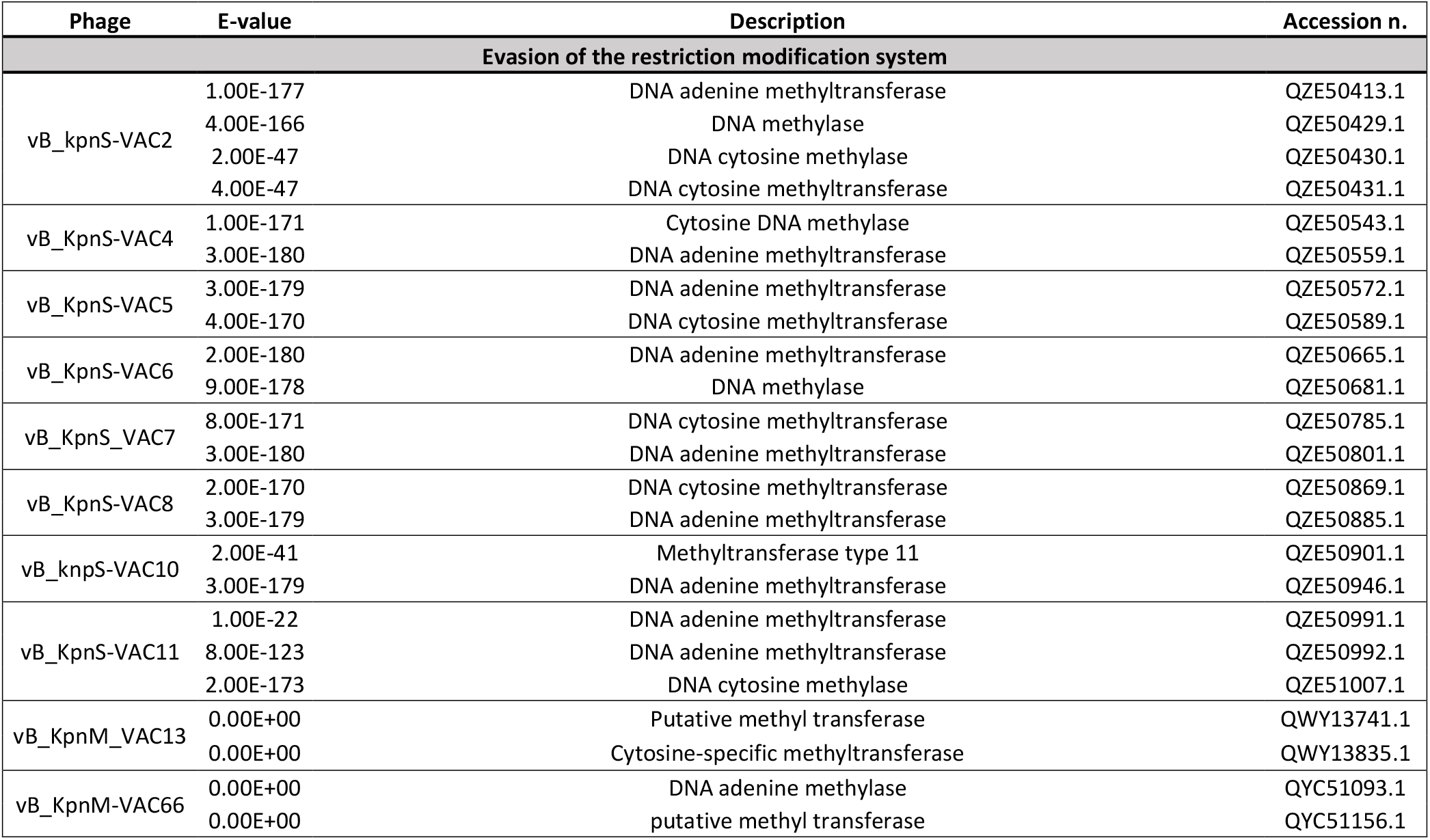
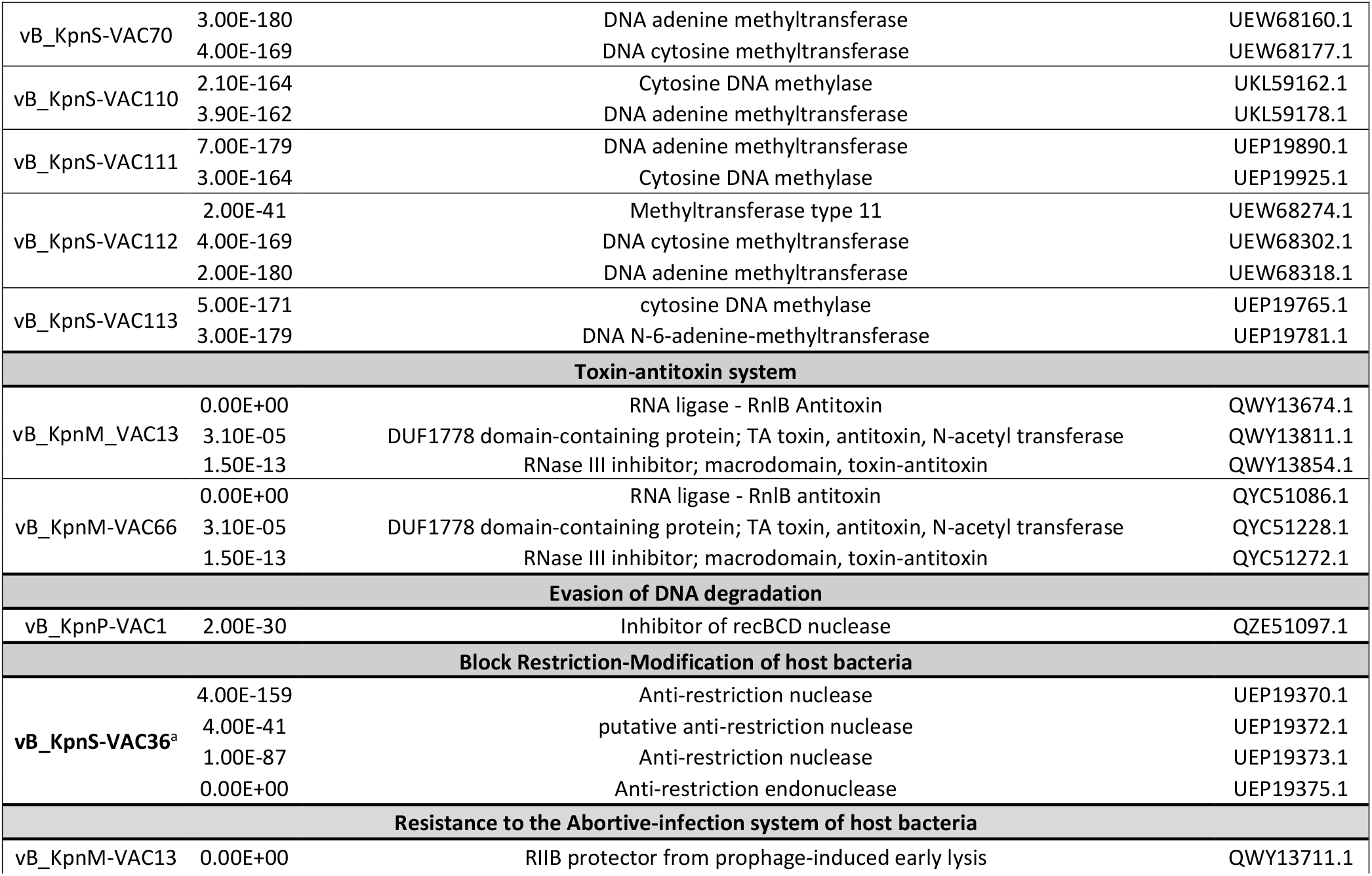
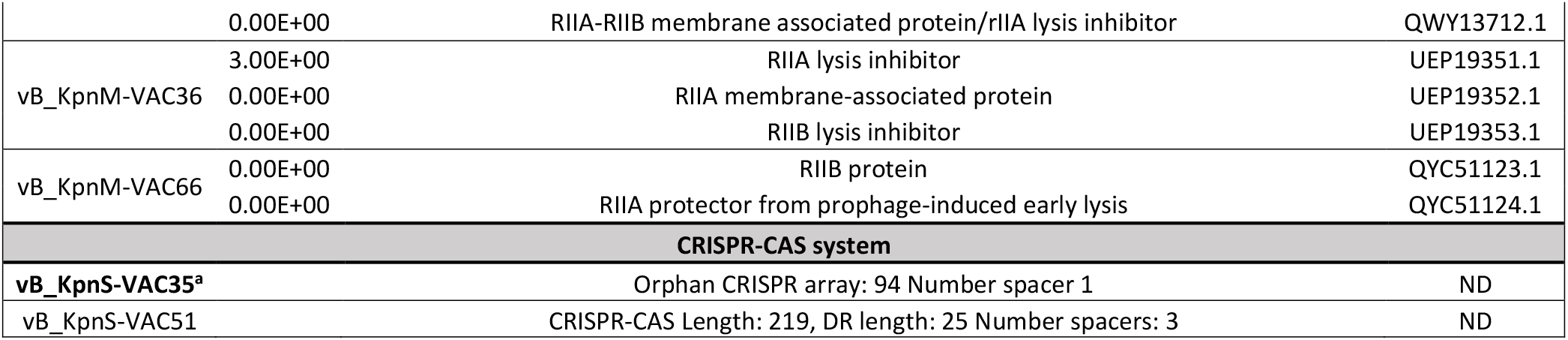
Phage defence mechanisms against bacteria detected in the phages under study. In bold, the clinical strains selected for phage interaction studies.

#### Phylogenetic analysis of phages

Phylogenetic analysis of the twenty-one phages revealed that they were clustered in the following families: i) *Drexlerviridae* (vB_KpnS-VAC2, vB_KpnS-VAC4, vB_KpnS-VAC5, vB_KpnS-VAC6, vB_KpnS-VAC7, vB_KpnS-VAC8, vB_KpnS-VAC10, vB_KpnS-VAC11, vB_KpnS-VAC35, vB_KpnS-VAC70, vB_KpnS-VAC110, vB_KpnS-VAC111, vB_KpnS-VAC112 and vB_KpnS-VAC113); ii) *Autographiviridae* (vB_KpnP-VAC1, vB_KpnP-VAC25 and vB_KpnP-VAC71); iii) *Ackerviridae* (vB_KpnS-VAC35 and vB_KpnS-VAC51); and iv) *Myoviridae* (vB_KpnM-VAC36, vB_KpnM-VAC13 and vB_KpnM-VAC66). It is worth mentioning that in the case of *Autographiviridae*, phage vB_KpnP-VAC1 was phylogenetically more similar to vB_KpnP-VAC71 than to B_KpnP-VAC25. Whereas, in the case of the *Myoviridae*, phage vB_KpnM-VAC36 was phylogenetically more distant than phages vB_KpnM-VAC13 and vB_KpnM-VAC66, which are very similar to each other, as demonstrated in a previous study conducted by our research group (38). (Figure 1A).

#### Genomic comparison

In the case of the *Drexilerviridae* family, the results show that phages vB_KpnS-VAC110 and vB_KpnS-VAC113 are very similar (query: 97 %, identity: 99.83 %), as are phages vB_KpnS-VAC7 and vB_KpnS-VAC4 (query: 91 %, identity: 98.93 %) and phages vB_KpnS-VAC111 and vB_KpnS-VAC112 (query: 90%, identity: 99.55 %). However, phages vB_KpnS-VAC70, vB_KpnS-VAC2, vB_KpnS-VAC6 and vB_KpnS-VAC111 show a low degree of similarity (query > 75%, identity: > 82 %). In the *Autographiviridae* family, a low degree of similarity between all phages was observed (query > 22 %, identity: > 76%). In the *Ackerviridae* family, partial similarity between phages vB_KpnS-VAC35 and vB_KpnS-VAC51 (query: 93 %, identity: 95.37 %) was observed. Finally, in the *Myoviridae* family, the phage vB_KpnM-VAC36 was very different from the other two phages, vB_KpnM-VAC13 (query: 0 %, identity: 80.75 %) and vB_KpnM-VAC66 (query: 3 %, identity: 85.05 %), which are very similar (query: 95 %, identity: 97.56%), as previously demonstrated in a study carried out by our research group (38) (Figure 2).

**Figure 2.**
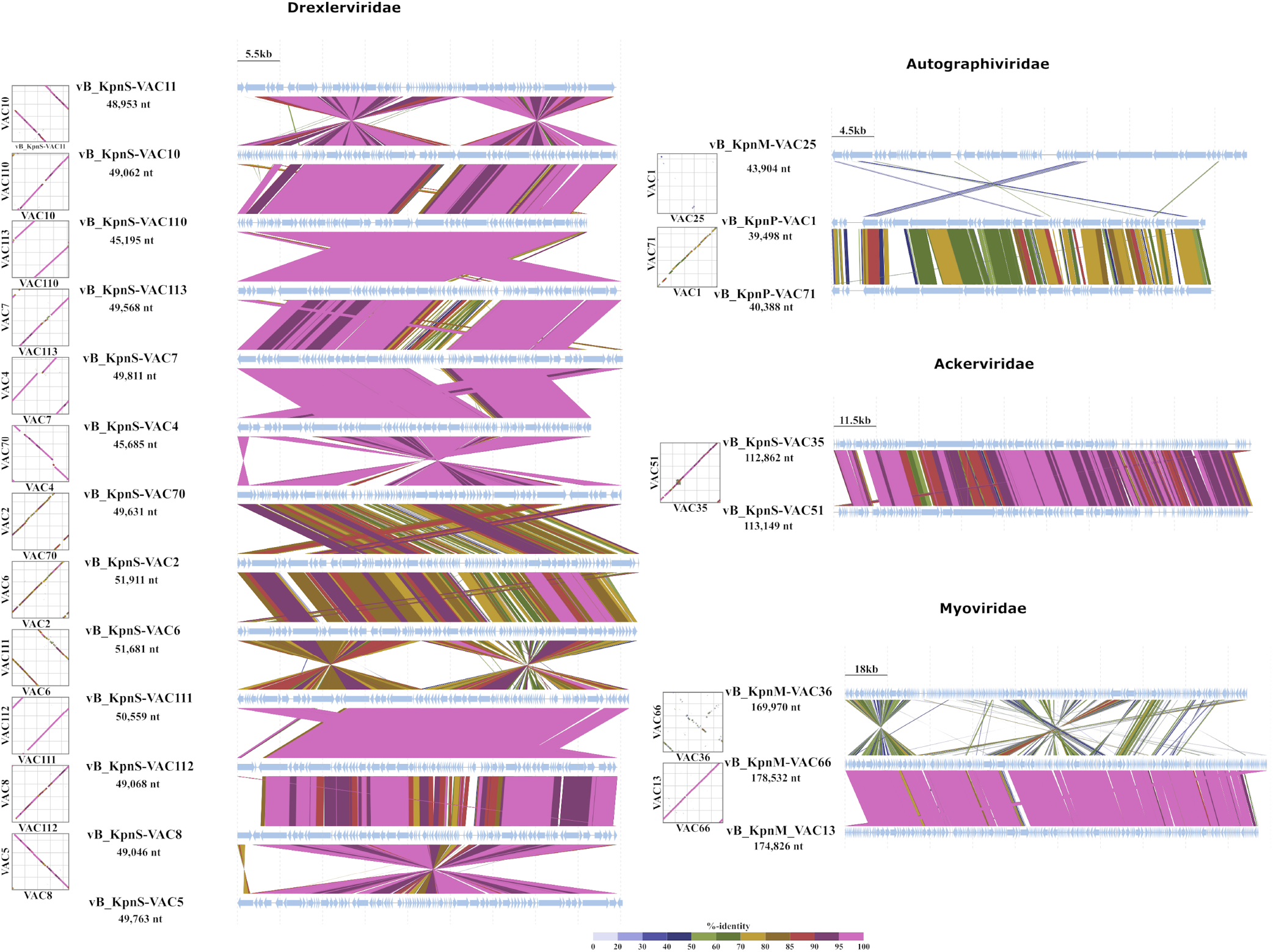
Graphic comparison of the homology of the twenty-one phages, grouped according to their families and in the same order as in the phylogenetic tree. The schematic representation was conducted with VipTree (https://www.genome.jp/viptree/, accessed in June 2022)

### Host-range assay

The phage infectivity in the collection of forty-seven clinical strains of *K. pneumoniae* is shown in Figure 3A. The results showed a high variability of infectivity between the phages (Figure 3B). Phage vB_KpnP-VAC1 had the lowest host-range, infecting only the strain K2986, while phage vB_KpnM-VAC13 presented the highest range of activity, infecting twenty-seven strains. Phages vB_KpnS-VAC35 and vB_KpnM-VAC36 also exhibited a high host-range, infecting both fourteen strains. Therefore, the following experiments focused on these two phages (vB_KpnS-VAC35 and vB_KpnM-VAC36).

**Figure 3.**
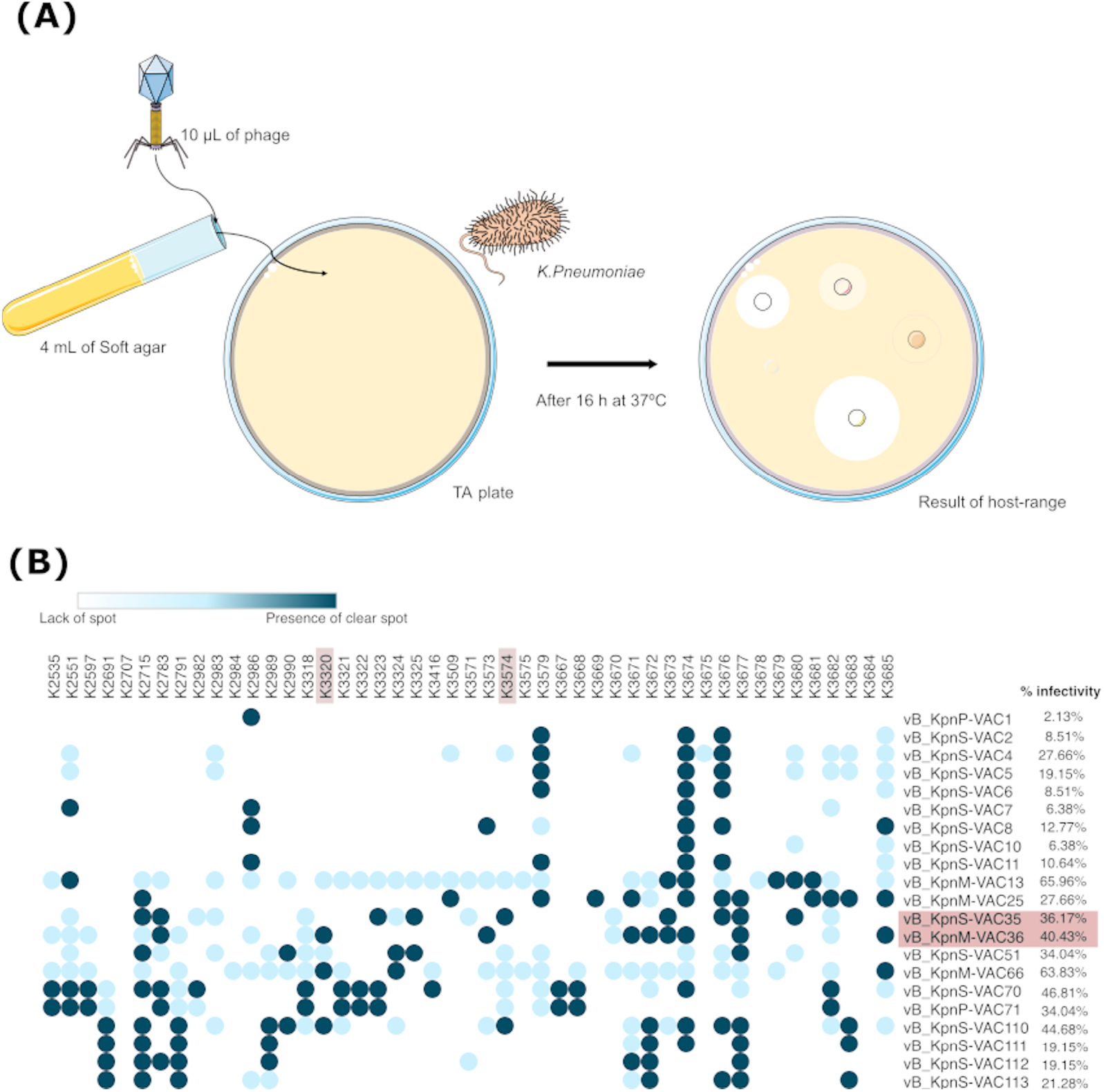
**(A)** Schematic representation of the host-range technique **(B)** Host-range of the twenty-one phages included in the collection of forty-seven clinical strains of *K. pneumoniae* and percentage of infectivity.

### Bacterial genome analysis: K3574 and K3320 clinical isolates

Both strains were found to have an intact CRISPR-Cas type I-E, with 35 spacers and 29 repeats in the case of K3574, and with 32 spacers and 29 repeats in the case of strain K3320 (Figure 4 A and B). In turn, five plasmids located in five different contigs was found in the strain K3474, while strain K3320 had four plasmids located in three different contigs (Table 4). On the other hand, only strain K3574 exhibited an RM system, which was type II and functioned as a methyltransferase (Table 4). Finally, five prophages (two intact and three questionable) were detected in strain K3574, and seven prophages (three intact, two incomplete and two questionable) were detected in strain K3320. However, only the data of the prophages considered intact are shown in Table 5.

**Figure 4.**
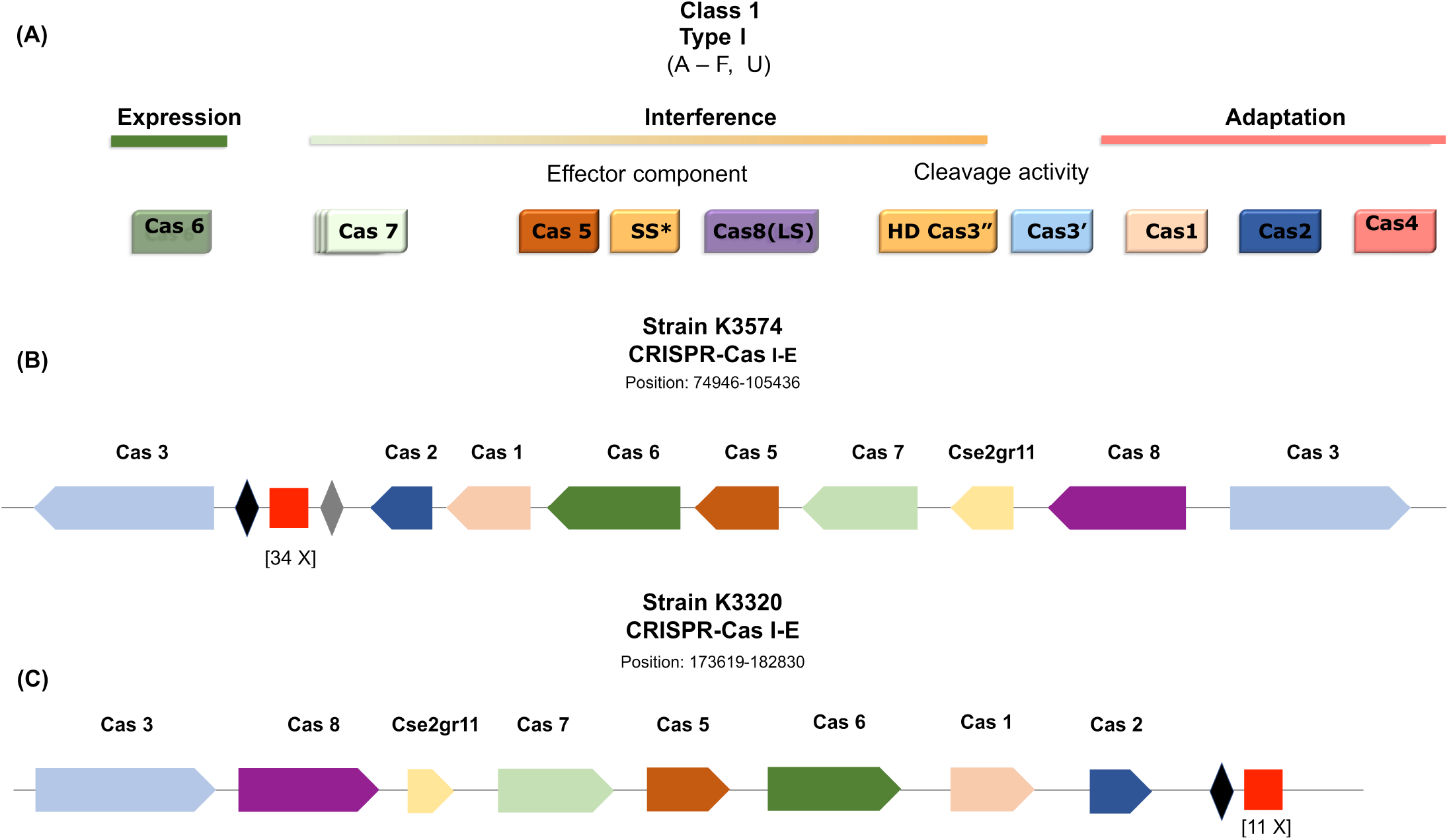
**(A)** Scheme of the modular organization of class I, type I CRISPR-Cas systems. Diagram adapted from Ishino *et al*. (2018). SS* indicates the putative small subunit (SS) that might be fused to the large subunit in several type I subtypes. (**B** and **C**) Graphic representation of the CRISPR-cas system of strains K3574 and K3320, and adapted image of CRISPR Miner 2 (http://www.microbiome-bigdata.com/CRISPRminer2/index/, accessed in March 2022).

**Table 4.** Plasmid and RM system of the selected clinical strains of *Klebsiella pneumoniae* K3574 and K3320.

| PLASMID |  |  |  |  |  |  |
| --- | --- | --- | --- | --- | --- | --- |
| Database | Plasmid | Identity | Contig | Start | Stop | Accession n. |
| K3574 |  |  |  |  |  |  |
| Enterobacterales | Col(pHAD28) | 97.78 | NODE_58_length_2240_cov_4.51213_ID_117 | 2148 | 2237 | KU674895 |
| Enterobacterales | Col440I | 96.49 | NODE_50_length_4243_cov_1686.76_ID_93 | 2628 | 2741 | CP023920 |
| Enterobacterales | IncFIA(HI1) | 98.97 | NODE_34_length_20733_cov_46.3618_ID_67 | 19450 | 19836 | AF250878 |
| Enterobacterales | IncFIB(K) | 98.93 | NODE_24_length_50018_cov_32.4049_ID_47 | 43034 | 43593 | JN233704 |
| Enterobacterales | IncFIB(pKPHS1) | 96.43 | NODE_1_length_470932_cov_21.7106_ID_1 | 156126 | 156685 | CP003223 |
| K3320 |  |  |  |  |  |  |
| Enterobacterales | Col(pHAD28) | 100.0 | NODE_100_length_2065_cov_2074.6_ID_199 | 1 | 100 | KU674895 |
| Enterobacterales | Col(pHAD28) | 100.0 | NODE_100_length_2065_cov_2074.6_ID_199 | 1886 | 2016 | KU674895 |
| Enterobacterales | IncFIB(K) | 98.93 | NODE_69_length_8285_cov_27.8857_ID_137 | 3503 | 4062 | JN233704 |
| Enterobacterales | IncR | 99.2 | NODE_76_length_6016_cov_46.5918_ID_151 | 648 | 898 | DQ449578 |
| RESTRICTION MODIFICATION SYSTEM |  |  |  |  |  |  |
| Type: Function | Gen: Recognition seq | Identity | Contig | Start | Stop | Recognition seq |
| K3574 |  |  |  |  |  |  |
| Type II R-M : Methyltransferase | M.Kpn34618DCM : CCWGG | 99.79 | Node_3_length_359415_cov_23.4099_ID_5 | 316287 | 317720 | CPO10392 |
| K3320 |  |  |  |  |  |  |
| NA | NA | NA | NA | NA | NA | NA |

**Table 5.**
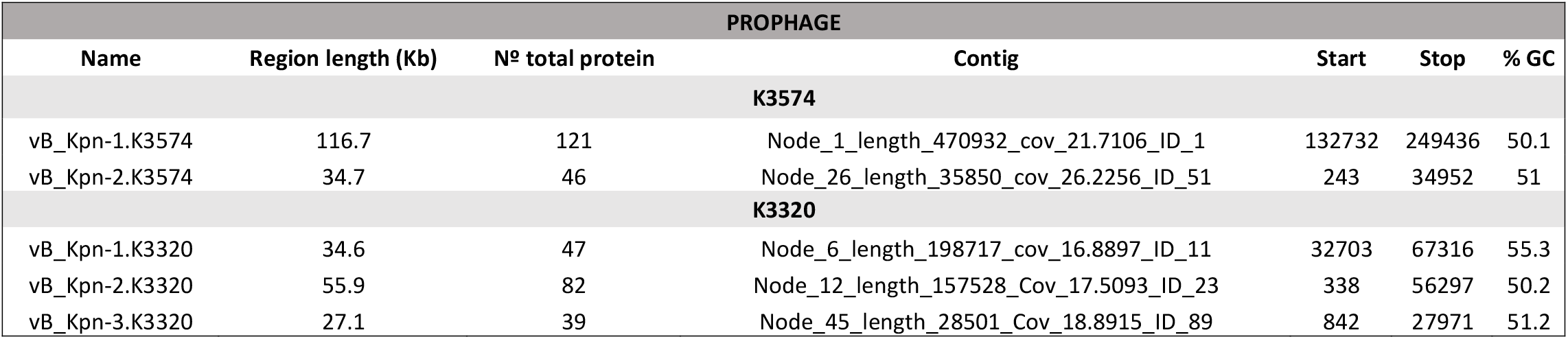
Intact prophage found in the genome of K3574 and K3320.

| PROPHAGE |  |  |  |  |  |  |
| --- | --- | --- | --- | --- | --- | --- |
| Name | Region length (Kb) | Nº total protein | Contig | Start | Stop | % GC |
| K3574 |  |  |  |  |  |  |
| vB_Kpn-1.K3574 | 116.7 | 121 | Node_1_length_470932_cov_21.7106_ID_1 | 132732 | 249436 | 50.1 |
| vB_Kpn-2.K3574 | 34.7 | 46 | Node_26_length_35850_cov_26.2256_ID_51 | 243 | 34952 | 51 |
| K3320 |  |  |  |  |  |  |
| vB_Kpn-1.K3320 | 34.6 | 47 | Node_6_length_198717_cov_16.8897_ID_11 | 32703 | 67316 | 55.3 |
| vB_Kpn-2.K3320 | 55.9 | 82 | Node_12_length_157528_Cov_17.5093_ID_23 | 338 | 56297 | 50.2 |
| vB_Kpn-3.K3320 | 27.1 | 39 | Node_45_length_28501_Cov_18.8915_ID_89 | 842 | 27971 | 51.2 |

### Characterization of phages vB_KpnS-VAC35 and vB_KpnM-VAC36

#### Phage adsorption

Adsorption of phages vB_KpnS-VAC35 and vB_KpnM-VAC36 (Figure 5 A and B) to the bacterial surface receptor was studied with the previously selected strains K3574 and K3320. Phage vB_KpnS-VAC35 showed a high percentage of adsorption, with 91.28 % of phage adsorbed in the strain K3574 after 5 min, while phage vB_KpnM-VAC36 showed slight adsorption in the strain K3320, with 39.02 % of phage adsorbed after 2 min (Figure 5 C and D).

**Figure 5.**
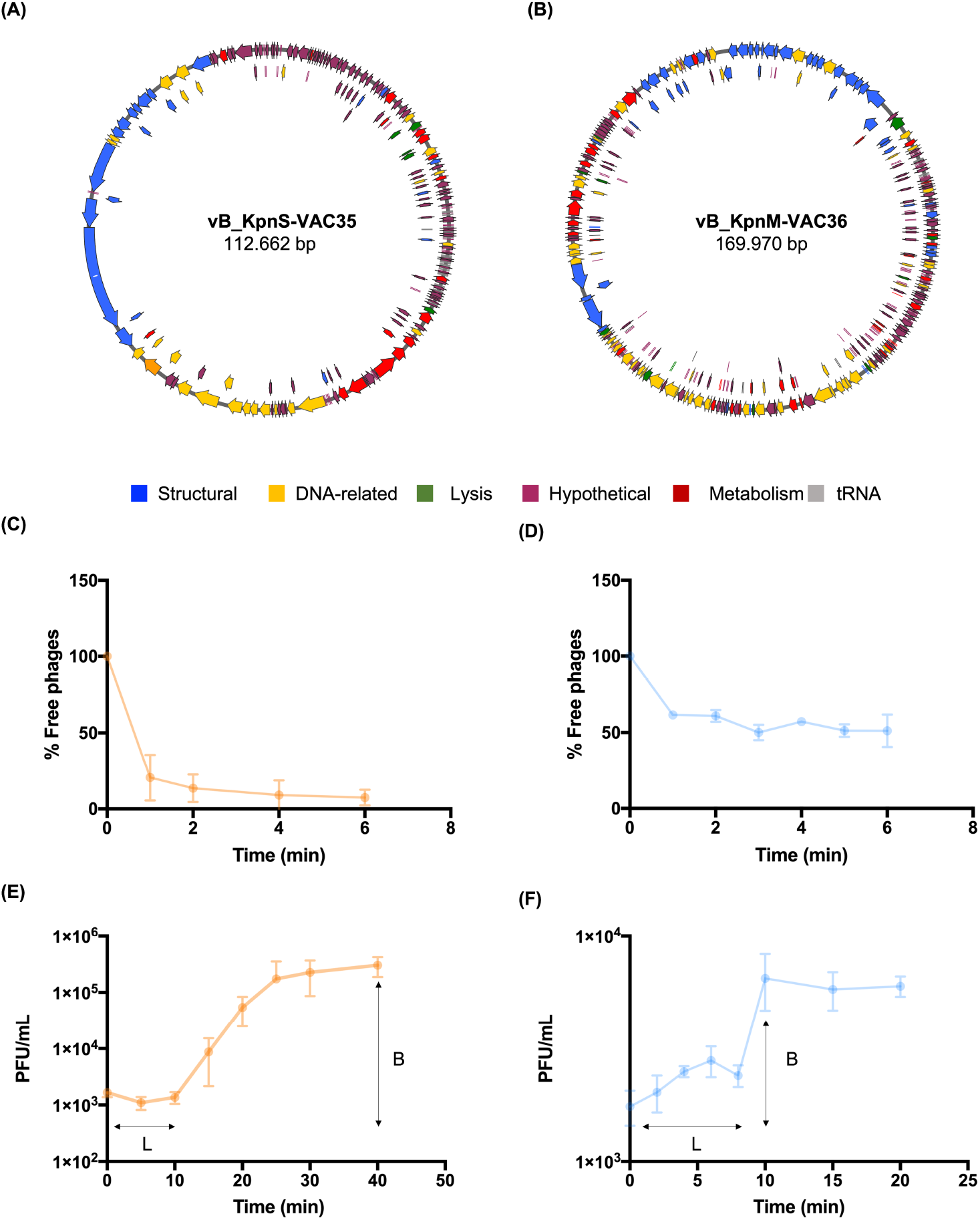
(**A** and **B**) Graphic representation of the genome of the phage vB_KpnS-VAC35 and vB_KpnM-VAC36, constructed with the Snapgene tool, version 6.0.5. **(C** and **D)** Adsorption curve for phages vB_KpnS-VAC35 and vB_KpnM-VAC36, with an adsorption time of 5 min and 2 min respectively. The error bar represents the standard deviation of the three experimental replicates. **(E** and **F)** One-step growth curve of phages vB_KpnS-VAC35 and vB_KpnS-VAC36, with a latent time (L) of 10 min and 8 min and a burst size (B) of 45.52 PFU/mL and 2.71 respectively. The error bar represents the standard deviation of the three experimental replicates.

#### One-step growth curve assay

The latent period determined by the one-step growth curve indicating the time taken for a phage particle to reproduce inside an infected host cell and the burst size (i.e. number of viral particles released in each infection cycle per cell) were respectively 10 min and 45.52 PFU/mL for phage vB_KpnS-VAC35 in strain K3574 and 8 min and 2.71 PFU/mL for the phage vB_KpnM-VAC36 in strain K3320 (Figure 5 E and F).

#### Phage kill curve of phage

The infectivity assay in liquid medium to determine the infection curve for phage vB_KpnS-VAC35 and vB_KpnS-VAC36 at a MOI of 1 in the selected clinical strains: K3574 and K3320, showed that both phages yielded successful infection, with OD_600nm_ values of respectively 0.155 ± 0.05 and 0.05 ± 0.01 reached after 1 h 30 of infection (Figure 6 A and B). In addition, the presence of resistant bacteria was not observed during the duration of the experiment (3 h). We monitored the CFU/mL and observed that for phage vB_KpnS-VAC35 in clinical strain K3574, the count reached 1.15 × 10^4^ ± 1.34 × 10^4^ CFU/mL. However, we observed a slight increase in the number of CFU/mL after 3 h of phage infection, reaching 2.35 × 10^4^ ± 2.12 × 10^4^ CFU/mL. Therefore, although the presence of resistant bacteria was not apparent from the optical density curves, the CFU/mL counts show that they were present. In phage vB_KpnM-VAC36 we observed a slight decrease in CFU/mL counts, which reached a value of 5.0 × 10^4^ ± 2.83 × 10^4^ after 1h 30 of phage infection and an increase of count after 2 h 30 of phage infection 1.35 × 10^5^ ± 3.54 × 10^4^ (Figure 6 C and D). Thus, although the appearance of resistant bacteria was not observed in the optical density test and the density remained unchanged, they did appear in the CFU/mL count test, as in the case of vB_KpnS-VAC35. Finally, we monitored the PFU/mL, and in both cases observed an increase in the number of PFU/mL after 30 min of phage infection, with values of 1.25 × 10^9^ ± 4.95 × 10^8^ PFU/mL for the phage vB_KpnS-VAC35 and 1.35 × 10^9^ ± 4.95 × 10^8^ PFU/mL for the phage vB_KpnM-VAC36 (Figure 5 E and F). Consequently, we can conclude that the number of CFU/mL is inversely proportional to the number of PFU/mL. These data confirm that the reduction in CFUs is due to multiplication of the phages.

**Figure 6.**
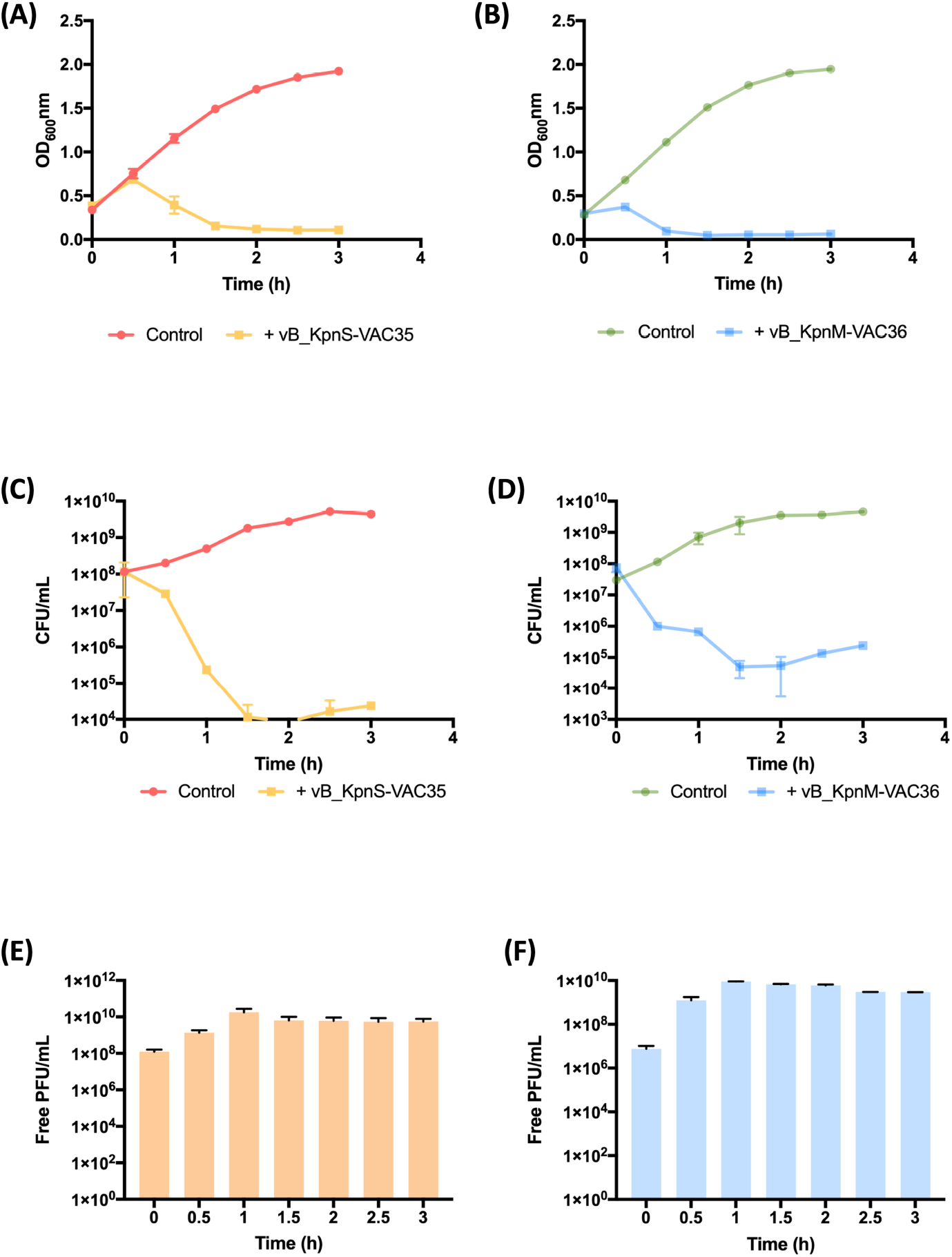
(**A** and **B**) Infection curve of the strains K3574 (orange) and K3320 (blue), respectively with phages vB_KpnS-VAC35 (light orange) and vB_KpnM-VAC36 (light blue) at an MOI of 1. **(C** and **D**) Measurement of viability by CFU/mL counts of strains K3574 and K3320 infected with respectively phage vB_KpnS-VAC35 and vB_KpnM-VAC36, at an MOI of 1, over time. **(E** and **F)** Measurement of PFU/mL counts of the phages vB_KpnS-VAC35 and vB_KpnM-VAC36 at an MOI 1 over time.

### NanoUHPLC-Tims-QTOF proteomic analysis: interaction between phages (vB_KpnS-VAC35 and vB_KpnM-VAC36) and clinical strains (K3574 and K3320)

The proteomic study conducted by NanoUHPLC-Tims-QTOF analysis revealed a large variety of proteins uniquely present in the phage-infected strain (listed in Figures 7 and 8 A, with the respective proportions in each strain): defence, resistance and virulence proteins, oxidative stress proteins, plasmid related proteins, cell wall related proteins and membrane proteins, as well as some transport proteins and proteins related to DNA, biosynthesis or degradation of proteins, ribosomes, metabolism, and some of unknown function. In the case of strain K3574 infected with phage vB_KpnS-VAC35, some prophage-related proteins were found, while in strain K3320 infected with phage vB_KpnM-VAC36 a large amount of tRNA was found. Regarding the defence proteins, we found proteins related to porins, multidrug efflux RND transporter, restriction-modification system type I methyltransferase, methyltransferase, two-component response regulator system, TA system type II RelE/ParE family, DNA starvation protein, fimbriae, pili (Figure 7 and 8 B). The details of all defence proteins, oxidative stress, plasmid/prophages and tRNA are summarised in Figures 7 and 8 B. The presence of some Acr candidates in the phage-infected strains was detected by NanoUHPLC-Tims-QTOF analysis: seven in K3574 infected by vB_KpnS-VAC35 and one in K3320 infected by the vB_KpnM-VAC36 phage (Table 6).

**Figure 7.**
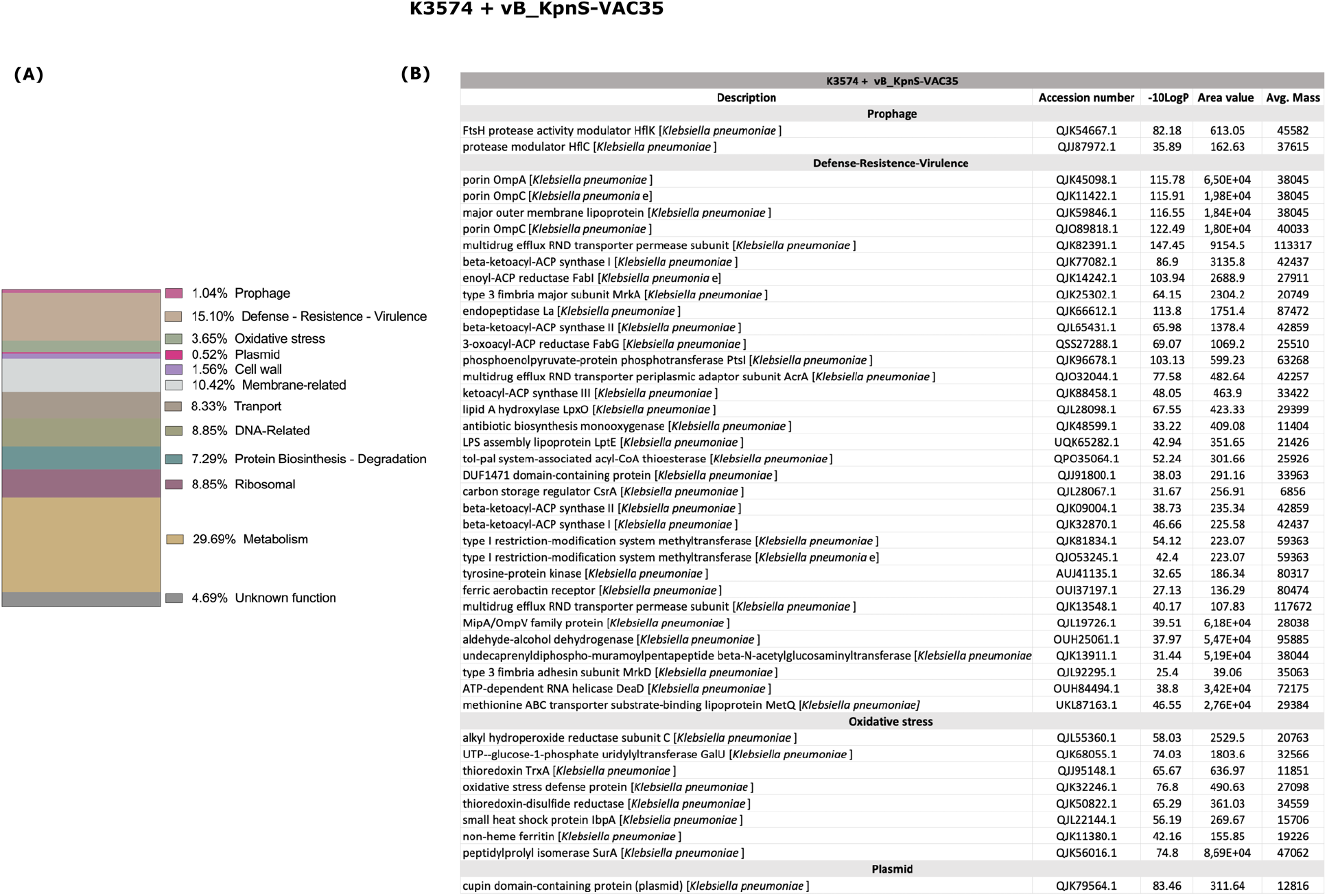
**(A)** Graphical representation of the proteomics results, showing the abundance of each group of proteins that are only present in the culture with the bacterial strain K3574 infected with phage vB_KpnS-VAC35. **(B)** Table showing the proteins of the functional groups of most relevance for phage-bacteria interactions: the other proteins present are listed in Supplementary Table 2. *Description* the protein header information as seen in the NCBI database, *-10LogP* the protein confidence score. *Area* the area under the curve of the peptide feature found at the same *m/z* and retention time as the MS/MS scan. This can be used as an indicator of the abundance and *Avg. Mass* is the protein mass calculated using the average mass.

**Figure 8.**
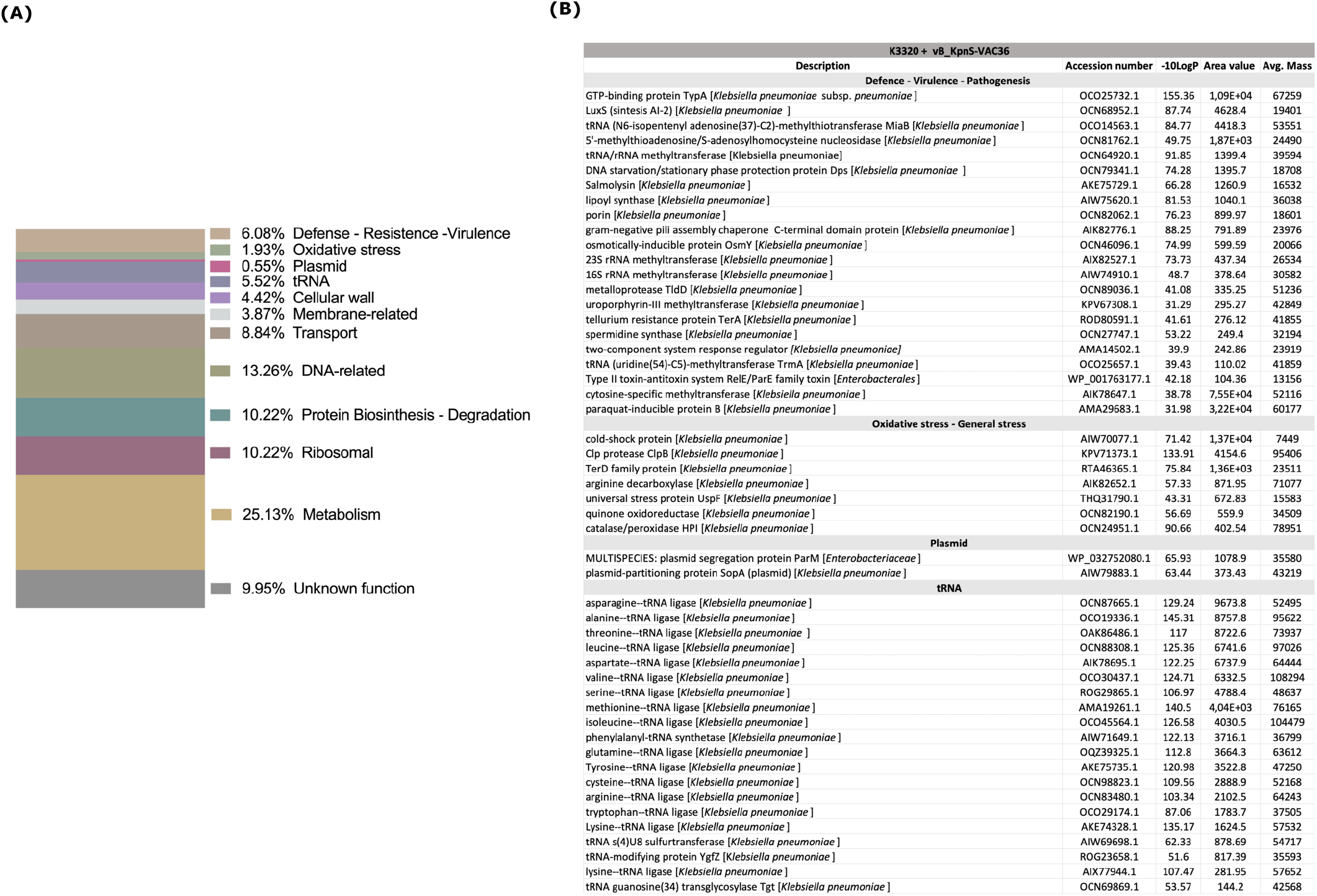
**(A)** Graphical representation of the proteomics results, showing the abundance of each group of proteins only present in the culture with the bacterial strain K3320 infected with phage vB_KpnM-VAC36. **(B)** Table showing the proteins of the functional groups of most relevance for phage-bacteria interaction: the other proteins present are listed in Supplementary Table 2. *Description* the protein header information as seen in the NCBI database, *-10LogP* the protein confidence score. *Area* the area under the curve of the peptide feature found at the same *m/z* and retention time as the MS/MS scan. This can be used as an indicator of the abundance and *Avg. Mass* is the protein mass calculated using the average mass.

**Table 6.** Acr candidate protein in phages vB_KpnS-VAC35 and vB_KpnM-VAC36 detected in the proteomic study (NanoUHPLC-Tims-QTOF). *Description* the protein header information as seen in the NCBI database, *Accession* the accession number of the protein as seen in the NCBI database, −*10LogP* the protein confidence score and *position* localization in the phage genome.

| Description | Accession number | -10LogP | Position |
| --- | --- | --- | --- |
| **K3574 + vB_KpnS-VAC35** |  |  |  |
| Hypothetical protein KPN4_89 (Acr candidate) [*Klebsiella* phage KPN4] | QEG11275.1 | 146.67 | 24497-24667 |
| 4-hydroxy-3-polyprenylbenzoate decarboxylase (Acr candidate) [*Klebsiella* phage vB_KpnS-VAC35] | UEP19035.1:3-122 | 107.65 | 22800-233162 |
| Hypothetical protein JIPhKp127_0059 (Acr candidate) [*Klebsiella* phage JIPh_Kp127] | QFR57489.1 | 104.09 | 23369-23911 |
| 6-phosphofructokinase (Acr candidate) [*Klebsiella* phage vB_KpnS-VAC35] | UEP19028.1 | 102.81 | 20466-20858 |
| Hypothetical protein (Acr candidate) [*Klebsiella* phage vB_KpnS-VAC35] | UEP19032.1 | 86.070 | 22064-22273 |
| Hypothetical protein (Acr candidate) [*Klebsiella* phage vB_KpnS-VAC35] | UEP19030.1:10-99 | 66.120 | 21405-21677 |
| Hypothetical protein (Acr candidate) [*Klebsiella* phage vB_KpnS-VAC35] | UEP19036.1 | 61.440 | 23155-23382 |
| **K3320 + vB_KpnM-VAC36** |  |  |  |
| Hypothetical protein (Acr candidate) [*Klebsiella* phage vB_KpnM-VAC36] | UEP19294.1 | 78.95 | 66295-66492 |

## DISCUSSION

Lytic phage therapy is currently considered one of the best alternatives for treating infections caused by multi-drug resistant bacterial pathogen (3, 4). Phages are known to exhibit some advantages over the use of antibiotics, including the continued warfare between phages and bacteria during the co-evolution of both organisms (12). Consequently, phages have developed defence mechanisms to evade the resistance mechanisms of bacteria (24–36), while at the same time bacteria have developed defence mechanisms to evade phage infection (46). In this context, the aims of the present study were to analyze twenty-one new lytic phages in search of defence mechanisms, and also to identify the defence mechanisms of two clinical strain K3574 and K3320 when infected by phages, as better knowledge of the latter will led to improvements in the use of phages to treat infections caused by MDR bacteria.

Regarding the results of the whole genome sequencing (WSG) and annotation, we observed that all of the phages belonged to the order *Caudoviridae*. Several studies have shown that dsDNA-tailed phages are the most abundant entity on earth (47, 48). Most of these phages are members of the *Siphoviridae* family (76.19 %), three are members of the *Myoviridae* family (14.29 %) and three are member of the *Podoviridae* family (14.29 %). Genome annotation has previously shown that all phages are lytic and lacking lysogenic genes such as integrase, recombinase and excisionase (20). This point is of vital importance for use of these phages in phage therapy (49, 50). Most phages were found to have a typical organization of the genome in functional modules, as previously described (2, 51, 52). By contrast, members of the *Myoviridae* family, which are included in the “larger phages” (> 100 bp), did not present specific lysis blocks, and structural and morphogenesis-related proteins were repeated in several blocks throughout the genome (38, 53). The genomes of all phages had endolysins and holins, proteins that are responsible for degradation of the bacterial cell wall during the infection by the host to facilitate the exit of the phage progeny (54).

Genomic annotation revealed the presence of numerous bacterial defence mechanisms: RM system evasion, TA system, DNA degradation evasion, blocking RM of host bacteria, genes that confer resistance to Abi system of host bacteria, a possible orphan CRISPR-cas system, and almost all the phages possessed a possible anti-CRISPR system. These mechanisms have all already been described (14). The anti-CRISPR, which is composed by operons of Acr and Aca proteins, was first discovered in 2013 in phages and prophages of *Pseudomonas aeruginosa* (55). Acr-Aca operons are defined as genomic loci fulfilling the following criteria: all genes should be in the same strand; all intergenic distances should be less than 150 bp; all genes encode proteins shorter than 200 amino acids in length; and finally at least one gene should be homologous to Acr or Aca proteins (56). The main problem of the search of new anti-CRIPSR is that Acr proteins are very poorly conserved, and the best way to discover new anti-CRISPR is therefore to use the guilt-by-association approach, which searches for Aca in the genome of phages. Although the function of Acas is not yet understood, these gene often encode a protein containing a helix-turn-helix (HTH) motif, suggesting that they fulfil a regulatory function (57).

The study of phage infectivity capacity revealed a large disparity in the infectivity, as previously demonstrated (58, 59): phages vB_KpnM-VAC13 and vB_KpnM-VAC66 displayed the highest infectivity capacity (38), whereas phage vB_KpnP-VAC1 displayed the lowest infectivity capacity (20). The wide host-range could be an advantage as it allows infection of a larger number of hosts (60), and this trait could be useful for successful phage therapy. In addition, the “larger phages” vB_KpnS-VAC35 (112.662 bp) and vB_KpnM-VAC36 (169.970 bp) showed a high capacity for infectivity, infecting fourteen clinical strains. Moreover, a possible anti-CRISPR system was detected in their genome. Thus, the presence of the CRISPR-Cas system in the bacterial strains that were successfully infected by these phages was examined to study the possible interaction of both defence mechanisms. The result of this search showed the presence of class I Type I-E intact CRISPR-Cas system in the genome of the strain K3574 and K3320. Both phages were examined with their respective host strains. The adsorption curve revealed that phage vB_KpnS-VAC35 displays a higher percentage of adsorption, a higher burst size and a longer latent period than phage vB_KpnM-VAC36. Moreover, analysis of the infectivity capacity by killing assay measuring the OD_600nm_, CFU/mL and PFU/mL revealed that phage vB_KpnS-VAC35 was more effective than phage vB_KpnM-VAC36.

Finally, proteomic studies were conducted with bacterial strains K3574 and K3320 with and without phage infection (vB_KpnS-VAC35 and vB_KpnS-VAC36) to determine any differences at the level of protein expression after phage infection. The pattern of protein expression was found to vary depending on the strain considered. This may be due to the different infection status of the bacterial cell at the time of sample processing or due to the inherent proprieties of the bacteria. Therefore, the results revealed the expression of FtsH protease modulator located in prophage of K3574 strain, which controls the lytic pathway (61), as well as overexpression of the cupin protein located in plasmid, a phosphomannose isomerase involved in LPS synthesis, which is an important determinant of pathogenicity and phage susceptibility (62). In addition, proteins related to bacterial defence, resistance and virulence and also to oxidative stress mechanisms (63) have been observed in the phage-infected strains. Overexpression of porins, efflux pumps, LPS and pili elements, previously described in the literature as phage receptors (64, 65), was also observed. Moreover, proteins involved in the *quorum* network were observed in both phage-infected strains, e.g. the LuxS that synthesizes AI-2 molecules, or the presence of the CsrA regulator in the strain K3320 infected by phage vB_KpnM-VAC36 (66, 67). Indeed, previous studies have associated the *quorum* network with phage infection (68, 69). In addition, a type II RelE/ParE TA system was expressed in strain K3320. This is a very interesting finding, as phage vB_KpnM-VAC36 did not successfully infect strain K3320. The fundamental role played by TA systems in the inhibition of phage infection has recently been demonstrated (20, 70–72). Interestingly, an inhibitor of the TA system (protein ID: QZE51102.1) was found in the genome of the phage vB_KpnP-VAC1. This type of gene, previously only described in one *E.coli* phage (33), may play a role in phage defence against bacteria.

The methyltransferases, other important proteins that play a key role in phage infection (73), were overexpressed in both K3574 and K3320 strains infected by phages. In addition, several Acr candidate proteins were expressed in the infected strains. This is a very interesting finding, because the anti-CRISPR could inhibit the host’s CRISPR-cas system and thus promote infection (57). Activation of all mechanisms could therefore be due to the phage-host interaction, with the bacteria trying to use all of their defence mechanisms in response to the infection. On the other hand, it was also observed that the protein biosynthesis and degradation machinery was highly activated, which may indicate the formation of new viral particles. Finally, a large amount of tRNA ligase was observed in strain K3320 infected with phage vB_KpnM-VAC36, whereas none was observed in strain K3574 infected with phage vB_KpnS-VAC35. This may be due to the presence of large amounts of tRNA in the genome of phage vB_KpnM-VAC36.

## CONCLUSION

Phage-host interactions have been examined ever since the discovery of phages a century ago. The present study revealed numerous defence mechanisms both against bacteria by phage (RM system evasion, TA system, DNA degradation evasion, RM block of host, resistance to Abi, anti-CRISPR and CRISPR-cas system) and against phage infection by bacteria (prophage, plasmid, defence/virulence/resistance and oxidative stress proteins). However, phage-host bacteria interactions remain poorly understood and further study is required in order to improve the efficacy of phage therapy.

## ACKNOWLEDGEMENTS

This study was funded by grant PI19/00878 awarded to M. Tomás within the State Plan for R+D+I 2013-2016 (National Plan for Scientific Research, Technological Development and Innovation 2008-2011) and co-financed by the ISCIII-Deputy General Directorate for Evaluation and Promotion of Research - European Regional Development Fund “A way of Making Europe” and Instituto de Salud Carlos III FEDER, Spanish Network for the Research in Infectious Diseases (REIPI, RD16/0016/0006 and CIBER21/13/00012, CB21/13/00084 and CIBER21/13/00095) and by the Study Group on Mechanisms of Action and Resistance to Antimicrobials, GEMARA (SEIMC, http://www.seimc.org/). M. Tomás was financially supported by the Miguel Servet Research Programme (SERGAS and ISCIII). I. Bleriot was financially supported by pFIS program (ISCIII, FI20/00302). O. Pacios L. Fernández-García and M. López were financially supported by a grant IN606A-2020/035, IN606B-2021/013 and IN606C-2022/002, respectively (GAIN, Xunta de Galicia). The authors acknowledge CESGA (www.cesga.es) in Santiago de Compostela, Spain for providing access to computing facilities and the RIAIDT-USC analytical facilities. Finally, we thank to researchers from the Spanish Network of Bacteriophages and Transducer Elements (FAGOMA) for contributing the lytic phages.

## AUTHOR CONTRIBUTIONS

### TRANSPARENCY DECLARATIONS

The authors declare that there are no conflicts of interest.

## SUPPLEMENTARY FILES

**Supplementary Table 1**. Possible Aca and Acr proteins of the anti-CRISPR system detected in the phages with the AcrDB bioinformatics tool (https://bcb.unl.edu/AcrFinder/, accessed in October 2021).

**Supplementary Table 2**. Table of the proteins found in the proteomic study by NanoUHPLC-Tims-QTOF in the strains infected with phages vB_KpnS-VAC35 and vB_KpnM-VAC36. *Description* the protein header information as seen in the NCBI database, *-10LogP* the protein confidence score. *Area* the area under the curve of the peptide feature found at the same *m/z* and retention time as the MS/MS scan. This can be used as an indicator of the abundance and *Avg. Mass* is the protein mass calculated using the average mass.

## REFERENCES

1. Bergh O, Børsheim KY, Bratbak G, Heldal M. High abundance of viruses found in aquatic environments. Nature. 1989;340(6233):467–8.

2. Bleriot I, Trastoy R, Blasco L, Fernández-Cuenca F, Ambroa A, Fernández-García L, et al. Genomic analysis of 40 prophages located in the genomes of 16 carbapenemase-producing clinical strains of Klebsiella pneumoniae. Microb Genom. 2020;6(5).

3. Domingo-Calap P, Delgado-Martínez J. Bacteriophages: Protagonists of a Post-Antibiotic Era. Antibiotics (Basel). 2018;7(3).

4. Taati Moghadam M, Khoshbayan A, Chegini Z, Farahani I, Shariati A. Bacteriophages, a New Therapeutic Solution for Inhibiting Multidrug-Resistant Bacteria Causing Wound Infection: Lesson from Animal Models and Clinical Trials. Drug Des Devel Ther. 2020;14:1867–83.

5. Furfaro LL, Payne MS, Chang BJ. Bacteriophage Therapy: Clinical Trials and Regulatory Hurdles. Front Cell Infect Microbiol. 2018;8:376.

6. Gordillo Altamirano FL, Barr JJ. Phage Therapy in the Postantibiotic Era. Clin Microbiol Rev. 2019;32(2).

7. Tkhilaishvili T, Merabishvili M, Pirnay JP, Starck C, Potapov E, Falk V, et al. Successful case of adjunctive intravenous bacteriophage therapy to treat left ventricular assist device infection. J Infect. 2021;83(3):e1–e3.

8. Lebeaux D, Merabishvili M, Caudron E, Lannoy D, Van Simaey L, Duyvejonck H, et al. A Case of Phage Therapy against Pandrug-Resistant Achromobacter xylosoxidans in a 12-year-old lung transplanted cystic fibrosis patient. Viruses. 2021;13(1).

9. Jennes S, Merabishvili M, Soentjens P, Pang KW, Rose T, Keersebilck E, et al. Use of bacteriophages in the treatment of colistin-only-sensitive Pseudomonas aeruginosa septicaemia in a patient with acute kidney injury-a case report. Crit Care. 2017;21(1):129.

10. Eskenazi A, Lood C, Wubbolts J, Hites M, Balarjishvili N, Leshkasheli L, et al. Combination of pre-adapted bacteriophage therapy and antibiotics for treatment of fracture-related infection due to pandrug-resistant Klebsiella pneumoniae. Nat Commun. 2022;13(1):302.

11. Principi N, Silvestri E, Esposito S. Advantages and Limitations of Bacteriophages for the Treatment of Bacterial Infections. Front Pharmacol. 2019;10:513.

12. Forterre P, Prangishvili D. The great billion-year war between ribosome-and capsid-encoding organisms (cells and viruses) as the major source of evolutionary novelties. Ann N Y Acad Sci. 2009;1178:65–77.

13. Labrie SJ, Samson JE, Moineau S. Bacteriophage resistance mechanisms. Nat Rev Microbiol. 2010;8(5):317–27.

14. Ambroa A, Blasco L, López M, Pacios O, Bleriot I, Fernández-García L, et al. Genomic Analysis of Molecular Bacterial Mechanisms of Resistance to Phage Infection. Front Microbiol. 2021;12:784949.

15. Lu MJ, Henning U. Superinfection exclusion by T-even-type coliphages. Trends Microbiol. 1994;2(4):137–9.

16. Lopatina A, Tal N, Sorek R. Abortive Infection: Bacterial Suicide as an Antiviral Immune Strategy. Annu Rev Virol. 2020;7(1):371–84.

17. Goldfarb T, Sberro H, Weinstock E, Cohen O, Doron S, Charpak-Amikam Y, et al. BREX is a novel phage resistance system widespread in microbial genomes. EMBO J. 2015;34(2):169–83.

18. Ofir G, Melamed S, Sberro H, Mukamel Z, Silverman S, Yaakov G, et al. DISARM is a widespread bacterial defence system with broad anti-phage activities. Nat Microbiol. 2018;3(1):90–8.

19. Song S, Wood TK. A Primary Physiological Role of Toxin/Antitoxin Systems Is Phage Inhibition. Front Microbiol. 2020;11:1895.

20. Bleriot I, Blasco L, Pacios O, Fernández-García L, Ambroa A, López M, et al. The role of PemIK (PemK/PemI) type II TA system from Klebsiella pneumoniae clinical strains in lytic phage infection. Sci Rep. 2022;12(1):4488.

21. Vasu K, Nagaraja V. Diverse functions of restriction-modification systems in addition to cellular defense. Microbiol Mol Biol Rev. 2013;77(1):53–72.

22. Song S, Wood TK. Post-segregational Killing and Phage Inhibition Are Not Mediated by Cell Death Through Toxin/Antitoxin Systems. Front Microbiol. 2018;9:814.

23. Guan L, Han Y, Zhu S, Lin J. Application of CRISPR-Cas system in gene therapy: Pre-clinical progress in animal model. DNA Repair (Amst). 2016;46:1–8.

24. Samson JE, Magadán AH, Sabri M, Moineau S. Revenge of the phages: defeating bacterial defences. Nat Rev Microbiol. 2013;11(10):675–87.

25. Meyer JR, Dobias DT, Weitz JS, Barrick JE, Quick RT, Lenski RE. Repeatability and contingency in the evolution of a key innovation in phage lambda. Science. 2012;335(6067):428–32.

26. Scholl D, Adhya S, Merril C. Escherichia coli K1’s capsule is a barrier to bacteriophage T7. Appl Environ Microbiol. 2005;71(8):4872–4.

27. Cornelissen A, Ceyssens PJ, Krylov VN, Noben JP, Volckaert G, Lavigne R. Identification of EPS-degrading activity within the tail spikes of the novel Pseudomonas putida phage AF. Virology. 2012;434(2):251–6.

28. Liu M, Deora R, Doulatov SR, Gingery M, Eiserling FA, Preston A, et al. Reverse transcriptase-mediated tropism switching in Bordetella bacteriophage. Science. 2002;295(5562):2091–4.

29. Labrie SJ, Tremblay DM, Moisan M, Villion M, Magadán AH, Campanacci V, et al. Involvement of the major capsid protein and two early-expressed phage genes in the activity of the lactococcal abortive infection mechanism AbiT. Appl Environ Microbiol. 2012;78(19):6890–9.

30. Krüger DH, Bickle TA. Bacteriophage survival: multiple mechanisms for avoiding the deoxyribonucleic acid restriction systems of their hosts. Microbiol Rev. 1983;47(3):345–60.

31. Warren RA. Modified bases in bacteriophage DNAs. Annu Rev Microbiol. 1980;34:137–58.

32. Iida S, Streiff MB, Bickle TA, Arber W. Two DNA antirestriction systems of bacteriophage P1, darA, and darB: characterization of darA-phages. Virology. 1987;157(1):156–66.

33. Sberro H, Leavitt A, Kiro R, Koh E, Peleg Y, Qimron U, et al. Discovery of functional toxin/antitoxin systems in bacteria by shotgun cloning. Mol Cell. 2013;50(1):136–48.

34. Otsuka Y, Yonesaki T. Dmd of bacteriophage T4 functions as an antitoxin against Escherichia coli LsoA and RnlA toxins. Mol Microbiol. 2012;83(4):669–81.

35. Deveau H, Barrangou R, Garneau JE, Labonté J, Fremaux C, Boyaval P, et al. Phage response to CRISPR-encoded resistance in Streptococcus thermophilus. J Bacteriol. 2008;190(4):1390–400.

36. Li Y, Bondy-Denomy J. Anti-CRISPRs go viral: The infection biology of CRISPR-Cas inhibitors. Cell Host Microbe. 2021;29(5):704–14.

37. Pacios O, Fernández-García L, Bleriot I, Blasco L, González-Bardanca M, López M, et al. Enhanced Antibacterial Activity of Repurposed Mitomycin C and Imipenem in Combination with the Lytic Phage vB_KpnM-VAC13 against Clinical Isolates of Klebsiella pneumoniae. Antimicrob Agents Chemother. 2021;65(9):e0090021.

38. Pacios O, Fernández-García L, Bleriot I, Blasco L, Ambroa A, López M, et al. Phenotypic and Genomic Comparison of Klebsiella pneumoniae lytic phages: vB_KpnM-VAC66 and vB_KpnM-VAC13. Viruses. 2021;14(1).

39. Wingett SW, Andrews S. FastQ Screen: A tool for multi-genome mapping and quality control. F1000Res. 2018;7:1338.

40. Ewels P, Magnusson M, Lundin S, Käller M. MultiQC: summarize analysis results for multiple tools and samples in a single report. Bioinformatics. 2016;32(19):3047–8.

41. Bankevich A, Nurk S, Antipov D, Gurevich AA, Dvorkin M, Kulikov AS, et al. SPAdes: a new genome assembly algorithm and its applications to single-cell sequencing. J Comput Biol. 2012;19(5):455–77.

42. Stamatakis A. RAxML version 8: a tool for phylogenetic analysis and post-analysis of large phylogenies. Bioinformatics. 2014;30(9):1312–3.

43. Kutter E. Phage host range and efficiency of plating. Methods Mol Biol. 2009;501:141–9.

44. Kropinski AM. Measurement of the rate of attachment of bacteriophage to cells. Methods Mol Biol. 2009;501:151–5.

45. Adriaenssens E, Brister JR. How to Name and Classify Your Phage: An Informal Guide. Viruses. 2017;9(4).

46. Rostøl JT, Marraffini L. (Ph)ighting Phages: How Bacteria Resist Their Parasites. Cell Host Microbe. 2019;25(2):184–94.

47. Ackermann HW. 5500 Phages examined in the electron microscope. Arch Virol. 2007;152(2):227–43.

48. Zinke M, Schröder GF, Lange A. Major tail proteins of bacteriophages of the order Caudovirales. J Biol Chem. 2022;298(1):101472.

49. Pirnay JP, Merabishvili M, Van Raemdonck H, De Vos D, Verbeken G. Bacteriophage Production in Compliance with Regulatory Requirements. Methods Mol Biol. 2018;1693:233–52.

50. Merabishvili M, Pirnay JP, De Vos D. Guidelines to Compose an Ideal Bacteriophage Cocktail. Methods Mol Biol. 2018;1693:99–110.

51. Casjens SR. Comparative genomics and evolution of the tailed-bacteriophages. Curr Opin Microbiol. 2005;8(4):451–8.

52. Fokine A, Rossmann MG. Molecular architecture of tailed double-stranded DNA phages. Bacteriophage. 2014;4(1):e28281.

53. Nogueira CL, Pires DP, Monteiro R, Santos SB, Carvalho CM. Exploitation of a Klebsiella bacteriophage receptor binding protein as a superior biorecognition molecule. ACS Infect Dis. 2021;7(11):3077–87.

54. Chen X, Liu M, Zhang P, Xu M, Yuan W, Bian L, et al. Phage-Derived Depolymerase as an Antibiotic Adjuvant Against Multidrug-Resistant. Front Microbiol. 2022;13:845500.

55. Bondy-Denomy J, Pawluk A, Maxwell KL, Davidson AR. Bacteriophage genes that inactivate the CRISPR/Cas bacterial immune system. Nature. 2013;493(7432):429–32.

56. Hang Le YB, Yi Haidong, Asif Amina, Wang Jiawi, Lighgow Trevor, Zhang Han, Ul Amir Afsar Minhas Fayyayz and Yin Yanbin. AcrDB: a database of anti-CRISPR operons in prokaryotes and viruses. Nucleic acids res. 2021; 49(D1):D622–629.

57. Trasanidou D, Gerós AS, Mohanraju P, Nieuwenweg AC, Nobrega FL, Staals RHJ. Keeping crispr in check: diverse mechanisms of phage-encoded anti-crisprs. FEMS Microbiol Lett. 2019;366(9).

58. Göller PC, Elsener T, Lorgé D, Radulovic N, Bernardi V, Naumann A, et al. Multi-species host range of staphylococcal phages isolated from wastewater. Nat Commun. 2021;12(1):6965.

59. Fong K, Tremblay DM, Delaquis P, Goodridge L, Levesque RC, Moineau S, et al. Diversity and Host Specificity Revealed by Biological Characterization and Whole Genome Sequencing of Bacteriophages Infecting. Viruses. 2019;11(9).

60. de Jonge PA, Nobrega FL, Brouns SJJ, Dutilh BE. Molecular and Evolutionary Determinants of Bacteriophage Host Range. Trends Microbiol. 2019;27(1):51–63.

61. Rokney A, Kobiler O, Amir A, Court DL, Stavans J, Adhya S, et al. Host responses influence on the induction of lambda prophage. Mol Microbiol. 2008;68(1):29–36.

62. Rahimi-Midani A, Kim MJ, Choi TJ. Identification of a Cupin Protein Gene Responsible for Pathogenicity, Phage Susceptibility and LPS Synthesis of Acidovorax citrulli. Plant Pathol J. 2021;37(6):555–65.

63. Sacher JC, Javed MA, Crippen CS, Butcher J, Flint A, Stintzi A, et al. Reduced Infection Efficiency of Phage NCTC 12673 on Non-Motile Campylobacter jejuni strains is related to oxidative stress. Viruses. 2021;13(10).

64. Maffei E, Shaidullina A, Burkolter M, Heyer Y, Estermann F, Druelle V, et al. Systematic exploration of Escherichia coli phage-host interactions with the BASEL phage collection. PLoS Biol. 2021;19(11):e3001424.

65. Stone E, Campbell K, Grant I, McAuliffe O. Understanding and Exploiting Phage-Host Interactions. Viruses. 2019;11(6).

66. Mey AR, Butz HA, Payne SM. Vibrio cholerae CsrA Regulates ToxR Levels in Response to Amino Acids and Is Essential for Virulence. mBio. 2015;6(4):e01064.

67. Ambroa A, Blasco L, López-Causapé C, Trastoy R, Fernandez-García L, Bleriot I, et al. Temperate Bacteriophages (Prophages) in Pseudomonas aeruginosa isolates belonging to the international cystic fibrosis clone (CC274). Front Microbiol. 2020;11:556706.

68. López M, Rueda A, Florido JP, Blasco L, Fernández-García L, Trastoy R, et al. Evolution of the Quorum network and the mobilome (plasmids and bacteriophages) in clinical strains of Acinetobacter baumannii during a decade. Sci Rep. 2018;8(1):2523.

69. Shah M, Taylor VL, Bona D, Tsao Y, Stanley SY, Pimentel-Elardo SM, et al. A phage-encoded anti-activator inhibits quorum sensing in Pseudomonas aeruginosa. Mol Cell. 2021;81(3):571–83.e6.

70. Song S, Wood TK. Toxin/Antitoxin System Paradigms: Toxins Bound to Antitoxins Are Not Likely Activated by Preferential Antitoxin Degradation. Adv Biosyst. 2020;4(3):e1900290.

71. Ni M, Lin J, Gu J, Lin S, He M, Guo Y. Antitoxin CrlA of CrlTA Toxin-Antitoxin System in a Clinical Isolate Pseudomonas aeruginosa inhibits lytic phage infection. Front Microbiol. 2022;13:892021.

72. LeRoux M, Srikant S, Teodoro GIC, Zhang T, Littlehale ML, Doron S, et al. The DarTG toxin-antitoxin system provides phage defence by ADP-ribosylating viral DNA. Nat Microbiol. 2022;7(7):1028–40.

73. Huang X, Wang J, Li J, Liu Y, Liu X, Li Z, et al. Prevalence of phase variable epigenetic invertons among host-associated bacteria. Nucleic Acids Res. 2020;48(20):11468–85.

74. Pacios O, Fernández-García L, Bleriot I, Blasco L, Ambroa A, López M, et al. Adaptation of clinical isolates of Klebsiella pneumoniae to the combination of niclosamide with the efflux pump inhibitor phenyl-arginine-β-naphthylamide (PaβN): co-resistance to antimicrobials. J Antimicrob Chemother. 2022;77(5):1272–81.

